# The ecology of Nipah virus in Bangladesh: a nexus of land use change and opportunistic feeding behavior in bats

**DOI:** 10.1101/2020.11.30.404582

**Authors:** Clifton D. McKee, Ausraful Islam, Stephen P. Luby, Henrik Salje, Peter J. Hudson, Raina K. Plowright, Emily S. Gurley

## Abstract

Nipah virus is a bat-borne paramyxovirus that produces yearly outbreaks of fatal encephalitis in Bangladesh. Understanding the ecological conditions that lead to spillover from bats to humans can assist in designing effective interventions. To investigate the current and historical processes that drive Nipah spillover in Bangladesh, we analyzed the relationship between spillover events and climatic conditions, the spatial distribution and size of *Pteropus medius* roosts, and patterns of land use change in Bangladesh over the last 300 years. We found that 53% of annual variation in winter spillovers is explained by winter temperature, which may affect bat behavior, physiology, and human risk behaviors. We infer from changes in forest cover that a progressive shift in bat roosting behavior occurred over hundreds of years, producing the current system where a majority of *P. medius* populations are small (median of 150 bats), occupy roost sites for 10 years or more, live in areas of high human population density, and opportunistically feed on cultivated food resources – conditions that promote viral spillover. Without interventions, continuing anthropogenic pressure on bat populations similar to what has occurred in Bangladesh could result in more regular spillovers of other bat viruses, including Hendra and Ebola viruses.

## Introduction

Zoonotic infections pose an increasing threat to human health [1,2], yet for many zoonoses we have a poor understanding of the biological factors that determine when and where animal hosts are infectious and pose a risk for spillover into human populations [3]. Spillover events often appear sporadic in space and time and repeated outbreaks are rare. This low replication makes it difficult to ascertain the natural history of pathogens. Moreover, rapid response to outbreaks of novel infectious diseases is facilitated when data on related pathogens have been collected through surveillance in animal hosts [4]. Only through long-term surveillance efforts that integrate knowledge of reservoir host ecology, routes of pathogen spillover, and the nature of human-animal interactions can we develop an understanding of the ecology of emerging infections and manage the risk of spillover [3]. Our goal in this study was to assess the ecological conditions that affect the spillover of Nipah virus from fruit bats to humans in Bangladesh based on almost two decades of outbreaks.

Nipah virus (family *Paramyxoviridae*, genus *Henipavirus*) is hosted by various *Pteropus* fruit bat species with partially overlapping ranges across countries of South and Southeast Asia [5–17] and potentially the Philippines, where an outbreak of illness in humans and horses from a Nipah-like virus occurred [18]. The range of henipaviruses including Hendra [19], Cedar [20], and others [21–23] extends throughout the geographic range of pteropodid bats to Australia, Indian Ocean islands, and sub-Saharan Africa [24]. These data, combined with limited evidence of pathology in henipavirus-infected bats [25,26], suggest that henipaviruses have had a long association with their bat reservoirs that spans the dispersal of pteropodid bats out of Southeast Asia to other regions [27–31].

Distinct outbreaks of Nipah virus infection have highlighted that the same pathogen may use multiple routes to spillover. Nipah virus was first discovered following an outbreak of febrile illness in pigs, pig farmers, and abattoir workers in Malaysia and neighboring Singapore between September 1998 and May 1999 [32–35]. The outbreak ended only after Malaysia established widespread surveillance of pigs, resulting in the culling of over one million animals [36]. Outbreaks of Nipah virus infection in Bangladesh have a very different ecological pattern. Since 2001 when the first cases of human encephalitis in Bangladesh and India were linked to Nipah virus [5,37], outbreaks have been reported almost every year in Bangladesh and more sporadically in neighboring India [38,39]. Outbreaks in Bangladesh are seasonal, with cases occurring between December and April [40] and cluster primarily in the central and northwest districts of the country. Unlike the outbreaks in Malaysia, those in Bangladesh do not involve an intermediate animal host and are instead linked to drinking fresh or fermented sap (*tari*) from silver date palm trees (*Phoenix sylvestris*) [41–43]. Geographic variation in observed spillover frequency across Bangladesh is partly explained by the proportion of households that drink fresh date palm sap [44] and the distance to the nearest hospital where systematic Nipah virus surveillance occurs [40]. The independence of these spillover events is supported by the genetic variability among Nipah virus sequences from humans and bats in Bangladesh collected from separate outbreaks, contrasting with the more homogeneous sequences from Malaysia [45]. Lastly, human-to-human transmission of Nipah virus occurs in Bangladesh [46,47] with an average reproduction number (the average number of secondary cases per case patient) of 0.33 (95% confidence interval [CI]: 0.19–0.59) estimated over 2001–2014 [47] or 0.2 (95% CI: 0.1–0.4) over 2007–2018 [38]. Human-to-human transmission of Nipah virus has also been reported during Nipah virus outbreaks in India in 2001, 2007, and 2018 [37,39,48,49]. Although human-to-human transmission was not widely acknowledged in Malaysia at the time of the outbreak [34], methods for detecting such transmission events (e.g., contact tracing) may not have been in place. Additionally, numerous cases reported in the literature had no contact with pigs, suggesting human-to-human transmission may be an alternative explanation [35,50,51]. Thus, the extent of human-to-human transmission that occurred during the Malaysian Nipah virus outbreak remains unclear.

One striking similarity between Nipah virus ecology in Bangladesh and Malaysia is that spillovers were facilitated by human resource supplementation in modified landscapes [52]. In Malaysia this involved planting fruit trees in close proximity to piggeries [53,54] whereas in Bangladesh the key resource appears to be date palm sap. *Pteropus medius* (formerly *P. giganteus*) frequently visit date palm trees to consume sap, potentially contaminating sap by licking the shaved area of the tree, urinating or defecating in the collection pots, or in some cases, becoming trapped and dying in the pot [42,55,56]. Visits by *P. medius* are highest during winter months (Islam et al., in preparation) when date palm sap is primarily harvested to drink fresh (October to March or April) [41,55,57] and when other available cultivated fruit resources for bats are low [58]. While *Phoenix sylvestris* is a native species in Bangladesh [59–62], date palm sap would not be available to bats if trees were not tapped by sap collectors. *P. medius* is found throughout Bangladesh and bats shed Nipah virus in their urine in all seasons [63]. Nipah virus can remain infectious at 22 °C in neutral pH bat urine for up to four days and artificial sap (13% sucrose, 0.21% bovine serum albumin, pH 7) for over one week [64,65]; most fresh sap and fermented *tari* is consumed within hours of collection [41,43,55]. While the prevalence of Nipah virus shedding in *P. medius* is generally low [63], presenting a bottleneck in spillover, the risk of foodborne transmission increases for communities with higher sap consumption [44].

These patterns imply that the spatiotemporal clustering of Nipah spillovers is a convergence of human and bat consumption behavior, wherein the risk of consuming sap contaminated with Nipah virus shed from bats is highest during winter when most sap is consumed by humans, and in regions with high rates of sap consumption.

However, there are still aspects of Nipah virus ecology in bats and their interface with human populations that are unclear. First, there is substantial year-to-year variation in the number of Nipah virus spillover events in Bangladesh [38] that may be explained by ecological factors influencing bat behavior and viral shedding. Cortes et al. [40] showed that differences in winter temperature can explain variation in Nipah virus spillovers, but this analysis only covered the period 2007–2013 and missed the decrease in spillovers observed after 2015 [38]. Second, we lack comprehensive information on the population biology, roosting and feeding behavior, and movement ecology of *P. medius* in Bangladesh. Like other *Pteropus* spp. bats, *P. medius* populations appear to be in decline due to hunting and habitat loss [66–68], but *P. medius* also appears to thrive in the human-dominated landscapes of Bangladesh. This adaptability derives from the opportunistic feeding habits of *Pteropus* species and their ability to forage over large areas [63,69–71]. Even though Bangladesh is already the most densely populated country that is not a small city-state or island [72], more *P. medius* roosts in Bangladesh are found in areas with higher human population density, forest fragmentation, and supplemental food resources from residential fruit trees [73,74]. However, villages with Nipah virus spillovers did not have more *P. medius* roosts or total bats in the village or within 5 km of the village boundary than villages where spillovers have not been detected [44]. National surveys of *P. medius* roost sites and population trends, including mapping of food resources used by bats, would provide a better understanding of *P. medius* interactions with humans. Lastly, we lack a historical perspective on how land use changes in Bangladesh may have influenced *P. medius* populations and behavior, thereby setting the stage for the emergence of Nipah virus. Analysis of these aspects of Nipah virus ecology will provide clearer insights into the potential drivers of Nipah virus spillover from bats.

The objective of this study was to describe the ecological factors that produce frequent spillover of Nipah virus, including climate effects on bat behavior or physiology, the geography of bat roosting sites in Bangladesh, and the relationship between historical land use change and bat roosting behavior. Following the results of Cortes et al. [40], we hypothesized that Nipah virus spillovers would have a strong relationship with winter temperature that explains annual variation in spillover numbers between 2001–2018. Regarding *P. medius* roosting sites, we hypothesized that spatial variables related to climate, human population density, land use, and anthropogenic food resources such as fruit trees and date palm trees could explain variation in the occupancy and size of roosting bat populations. Finally, we hypothesized that land use change, specifically the loss of primary forests, has been a continuous process throughout human occupation of the region that was accelerated during British occupation. This progressive loss of forests likely led to a shift in roosting sites toward more urban areas closer to anthropogenic food resources, a condition that facilitates spillover but predates the first recognized outbreaks of Nipah virus infection by many years. By assessing these patterns, we develop a more comprehensive view of Nipah virus ecology in Bangladesh and provide a path forward for research and management of this system.

## Materials and Methods

### Nipah virus spillover events

To investigate the spatial and temporal patterns of Nipah virus spillover in Bangladesh, we compiled data on the number of spillover events and affected administrative districts during 2001–2018. Cases prior to 2007 were detected through community investigations following reports of clusters of encephalitis. Cases from 2007 onward reflect those identified through systematic surveillance for Nipah virus infection at three tertiary care hospitals combined with investigations of all cases detected to look for clusters, as well as any reports of possible outbreaks through media or other information sources [38]. Independent spillover events were defined as index cases of Nipah virus infection within a given outbreak year. This definition excludes cases that resulted from secondary human-to-human transmission following spillover.

### Climate data

Expanding on the results from Cortes et al. [40] showing associations between climate and the number of spillover events during 2007–2013, we used data from 20 weather stations in Bangladesh. Mean temperature at three-hour intervals and daily precipitation between 1953–2015 were obtained from the Bangladesh Meteorological Department. Daily temperature and precipitation summary data from 2015 onwards were obtained from the National Climatic Data Center [75] and merged with the older data. We also downloaded monthly indices for three major climate cycles that lead to temperature and precipitation anomalies in the region: the multivariate ENSO index (MEI) for the El Niño–Southern Oscillation, the Indian Ocean dipole mode index (DMI), and the subtropical Indian Ocean dipole index (SIOD). Data were retrieved from the Japan Agency for Marine-Earth Science and Technology Application Laboratory [76] and the National Oceanic and Atmospheric Administration Physical Sciences Laboratory [77]. Based on the frequency of Nipah virus spillovers occurring in winter, we focused on weather summary statistics for each year that covered the period from the start of the preceding December to the end of February of a focal outbreak year. We calculated the mean and recorded the minimum temperature over all stations, the percentage of days below 17 °C, and the cumulative precipitation from all stations over the focal period. The choice of 17 °C was arbitrary but represents an upper bound for relative coolness during winter that does not produce any zeros. Mean winter MEI, DMI, and SIOD values were also calculated for each year.

### Survey of bat roost sites and food resources

The spatial distribution of *Pteropus medius* in Bangladesh was inferred from a country-wide survey of villages as part of investigations regarding risk factors for Nipah spillover performed over the winters of 2011–2012 and 2012–2013 [44]. Briefly, trained teams of data collectors interviewed key informants within villages, who identified known bat roost sites (both occupied and unoccupied) in the village and within 5 km of the village and reported details of the duration of roost occupancy and perceived population trends. The interviewers also mapped the location and number of date palm trees (*Phoenix sylvestris*) and known feeding sites that bats were reported to visit within 500 m of the villages. Feeding sites included fruit trees planted in orchards or in residential areas: jujube (*Ziziphus mauritiana*), banana, mango, guava, lychee, star fruit, jackfruit, papaya, sapodilla (*Manilkara zapota*), mulberry, hog plum (*Spondias mombin*), Indian olive (*Elaeocarpus serratus*), and other species.

### Spatial covariates of bat roost sites

To evaluate spatial covariates that could explain the occupancy (presence/absence of bats) and abundance (estimated population size) of bats living in mapped roost sites, we extracted data from available raster surfaces describing human population density, land use, bioclimatic variables (e.g., mean annual temperature and precipitation), elevation, slope, and forest cover. Spatial covariate raster files were downloaded from WorldPop [78,79], the Socioeconomic Data and Applications Center (SEDAC) [80], WorldClim [81], and a study on global forest cover change [82]. We also calculated the distance from an index roost site to the nearest village, neighboring roost, date palm tree, and feeding site, and the number of villages, other mapped roosts, date palm trees, and feeding sites within a 15 km radius around each roost. Average nightly foraging distances of individual *P. medius* in two colonies in Bangladesh were estimated to be 10.8 km and 18.7 km, so 15 km was chosen to represent the distance a bat might expect to travel to reach a suitable feeding site [63]. The number of potential covariates was initially reduced by removing variables that were colinear (Pearson’s correlation greater than 0.7). Descriptions, sources, spatial resolution, and distribution statistics for all 32 covariates are provided in Table A1.

### Historical land use data

Given the reliance of *P. medius* on tall trees for roosting and various native and cultivated fruit trees for food, we gathered data on historical changes in land use, particularly forested lands, across Bangladesh from data sources covering separate but overlapping time periods. Reconstructed natural biomes and anthropogenic biomes from 1700–2000 were extracted from rasters produced by Ellis et al. [83] using the HYDE 3.1 data model [84] and available from SEDAC. We reclassified their land use subcategories into three primary categories: dense settlements, consisting of urban and suburban areas with high human population density (>100 persons/km^2^ for settlements, >2500 persons/km^2^ for urban areas); rice villages and other croplands or rangelands; and forested areas, including populated woodlands and remote forests. Land use data for the years 1992, 2004, 2015, and 2018 were downloaded from the Organisation for Economic Co-operation and Development (OECD) land cover database [85], derived from European Space Agency Climate Change Initiative Land Cover maps [86]. Data for 1990 and 2016 were provided by the World Bank [87]. Land cover over the period 1930–2014 came from an analysis by Reddy et al. [88]. Finally, forest cover from 2000 and subsequent forest loss as of 2017 were calculated from maps produced by Hansen et al. [82] using the R package *gfcanalysis* [89,90]. For the calculations from Hansen et al. data, we chose a cutoff of 40% forest cover density to match the definition of dense forests used by Reddy et al. Across these datasets, we calculated the percentage of Bangladesh’s total land area (147,570 km^2^ [88]) that was classified as forest.

### Statistical analysis

Separate Nipah virus spillover events were clustered geographically by the latitude and longitude of affected administrative districts and temporally by the date of illness of each index case using a bivariate normal kernel via the R package *MASS* [91]. To examine the association between Nipah virus spillovers and climate variables, separate generalized linear models were produced that examined climate summary statistics and the number of spillover districts or independent spillover events assuming a Poisson distribution for each response. Model selection was performed to choose the best fitting combination of climate covariates according to Akaike’s information criterion corrected for small sample sizes (AICc) [92] using the R package *MuMIn* [93].

The importance of spatial covariates in explaining variation in the occupancy and abundance of bats at roost sites was assessed through a combination of linear modeling and machine learning. The covariates were standardized, and data were split into two sets: an occupancy dataset of 488 mapped roost sites with a binary variable describing whether bats were currently present or not and an abundance dataset of 323 mapped roost sites with the estimated count of bats at each currently occupied roost at the time of the interview. Both datasets were split into training (80%) and testing (20%) sets for validation of models [94]. Generalized linear models (GLMs) were fit with all potential covariates, assuming a binomial distribution for roost site occupancy and a negative binomial distribution for roost counts, which was chosen because of the observed overdispersion of the data, with a variance:mean ratio greater than unity. Due to the large number of potential covariates, least absolute shrinkage and selection operator (lasso) regularization was implemented to reduce the number of covariates and minimize prediction error [95]. We also used random forests to perform covariate selection and assess explanatory power [96]. This machine learning method constructs many decision trees using random subsets of the response variable and covariates then averages the predictions. This method of constructing and averaging a set of uncorrelated decision trees reduces overfitting relative to single decision trees. Linear modeling and random forests were performed in R using the packages *caret, glmnet*, and *ranger* [97–99].

## Results

### Spatiotemporal patterns of Nipah virus spillover

Based on 183 spillover events from 2001–2018, we confirmed previous analyses [38,40,44] showing that Nipah virus spillovers are spatially clustered within districts in the central and northwest regions of Bangladesh (Figure 1A). Outbreak years vary in the intensity of spillover and winter is the primary season when spillovers occur throughout the country (Figure 1B,C), although there are occasional events in early spring in central Bangladesh. With the exception of 2002, 2006, and 2016, Nipah virus spillovers have been observed every year since the virus was first identified in 2001, and as observed by Nikolay et al. [38], more spillovers were observed between 2010–2015 than before or after this period (Figure 1D). In accordance with previous work [40] covering 2007–2013, we confirmed that much of this yearly variation in spillover events (53%) can be explained by winter weather over the longer period 2001–2018. Mean winter temperature, minimum winter temperature, and the percentage of days below 17 °C all showed statistically significant associations with yearly spillover events and the number of affected districts (P < 0.001; Figures A1–A3). There were no significant associations with cumulative winter precipitation (P > 0.05; Figure A4) or the three climate oscillation indices (MEI, DMI, and SIOD; Figure A5). The percentage of days below 17 °C was chosen as the single best fitting covariate for both outcomes according to AICc (Tables A2–A3), showing that colder winter temperatures were associated with more spillovers and more affected districts during 2010–2015, followed by fewer spillovers and affected districts during the relatively warmer period of 2016–2018 (Figure 1D,E; Figure A3). Sensitivity analysis of the association between spillovers and the number of winter days below a certain temperature threshold confirmed that the relationship was strongest at thresholds of 16 to 18 °C, but was statistically significant for thresholds ranging from 15 to 20 °C. We note that spillover observations prior to 2007 mostly appear as undercounts relative to those expected by the winter temperatures (Figure 1E; Figures A1–A3), which may be attributed to the lack of systematic surveillance during that period [38].

**Figure 1.**
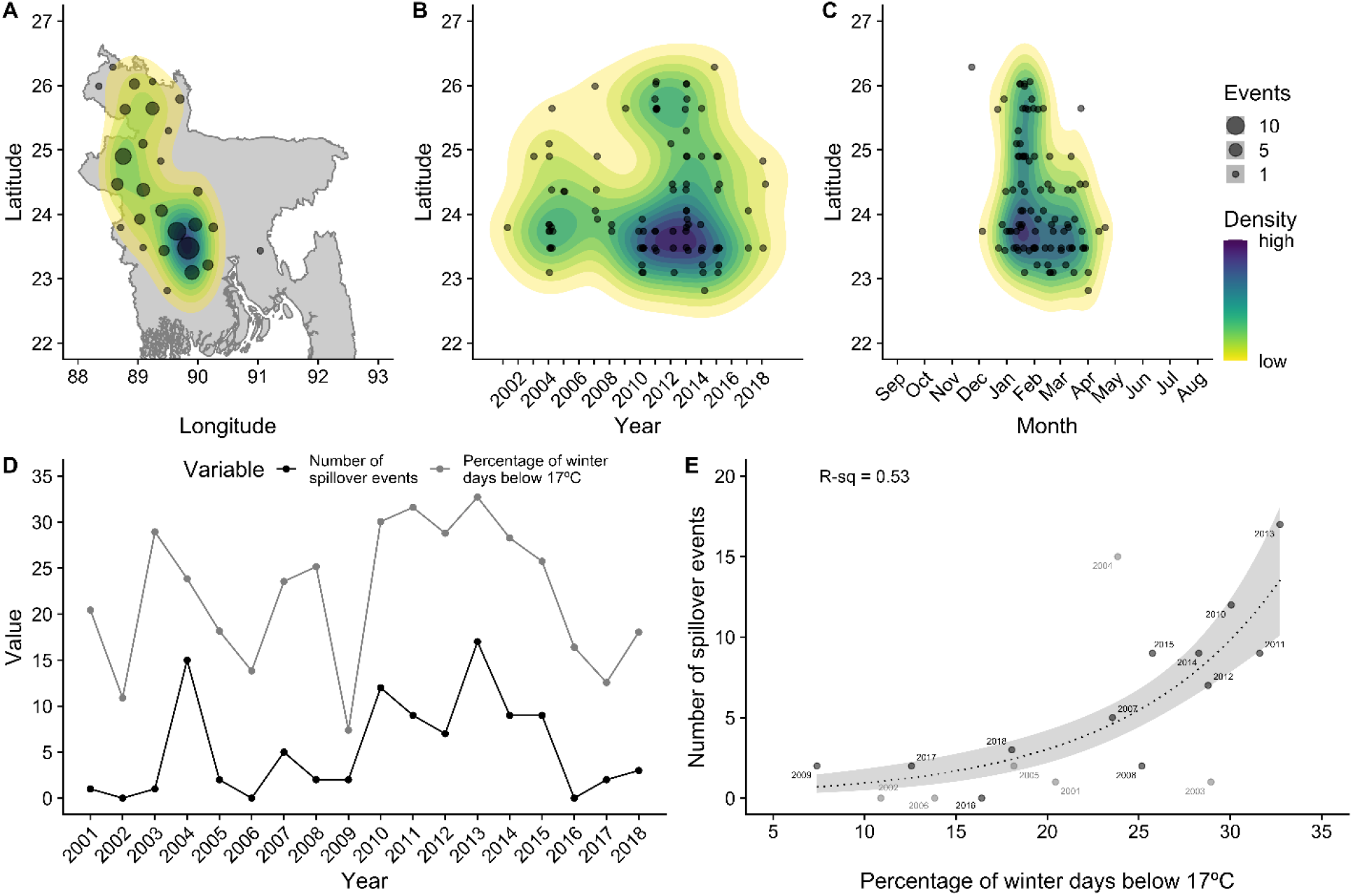
Spatiotemporal patterns of Nipah virus spillover events across Bangladesh, 2001–2018. Color contours in panels A–C show the spatial density of events estimated with a bivariate normal kernel. Panels D–E show the variation in the number of Nipah spillover events across years and the association with cold winter temperatures. Gray dots in panel E show the years before systematic Nipah virus surveillance.

### *Spatial distribution and sizes of* Pteropus medius *roosts*

Interviewers mapped a total of 474 roost sites in and around 204 villages, 315 that were occupied at the time of the interview and 159 that were unoccupied. According to interviewees, most occupied roosts (186, 59%) were reported as being at least occasionally occupied for more than 10 years, with an average occupancy duration of 8.5 years (Figure 2A). The majority (294, 93%) of roosts were reported to be continuously occupied every month within the last year, with an average duration of 11.6 months (Figure 2B). This pattern of continuous occupancy was reported by interviewees to have been similar over the last 10 years (Figure 2C). Interviewees generally could not recall what season bats began roosting at sites, but when reported, roosts were first occupied more frequently in winter than other seasons (Figure A6A). When considering intermittently occupied roost sites (<12 months of occupancy in a year), bats were also more likely to be present at roost sites during winter (Figure A6B).

**Figure 2.**
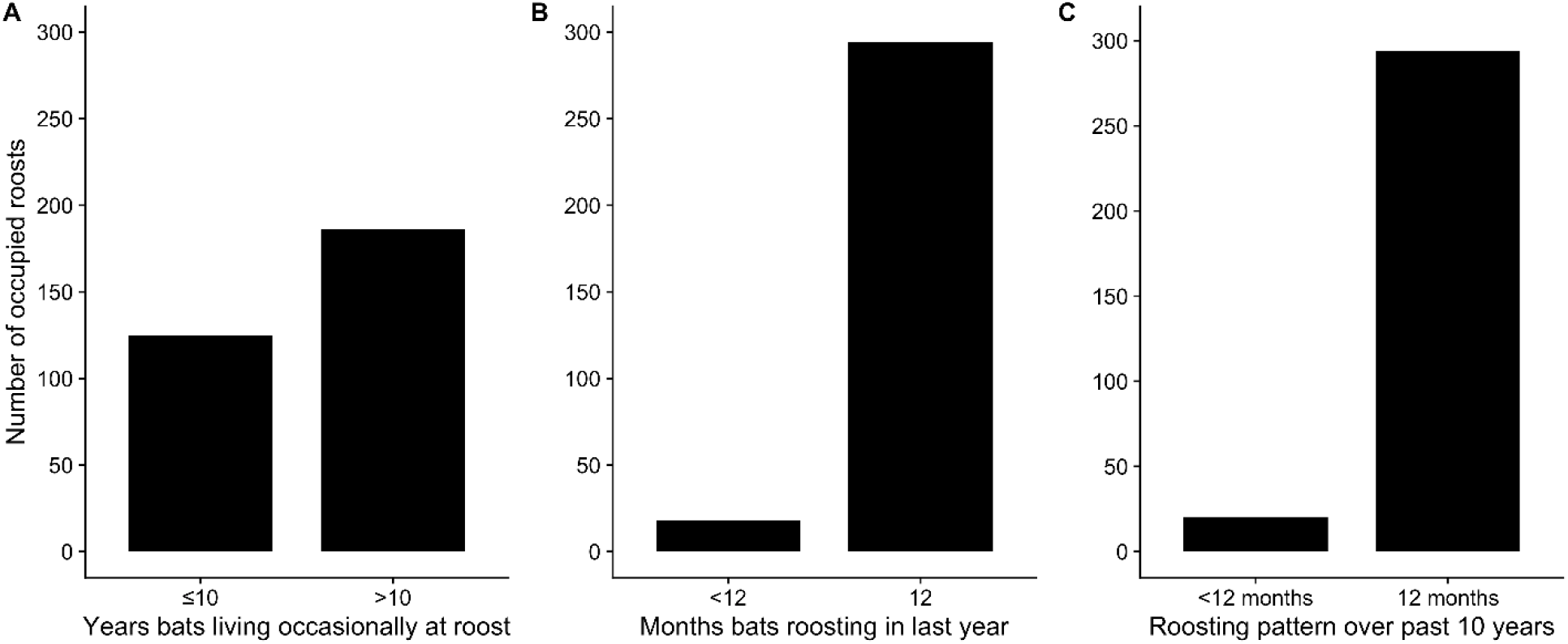
Duration of occupancy of *Pteropus medius* populations at occupied roost sites. According to interviewees, occupied roosts were most frequently occupied for more than 10 years (A) and for 12 months out of the year (B). Continuous roost occupancy was similar over the past 10 years (C).

The size of occupied roosts varied widely, from only one bat to an estimated 8,000 bats at one roost in west-central Bangladesh, with a median size of 150 bats (Figure 3A,B). Studies of *P. medius* demonstrate that this distribution of individual roost sizes is similar to those reported in Pakistan, India, Nepal, and Sri Lanka [100–106]. This contrasts with reports of much larger roosts of thousands of *P. lylei* in Cambodia and Thailand [13,107], and roost sizes of *P. alecto* and *P. poliocephalus* in Australia estimated in the tens of thousands [108–110].

**Figure 3.**
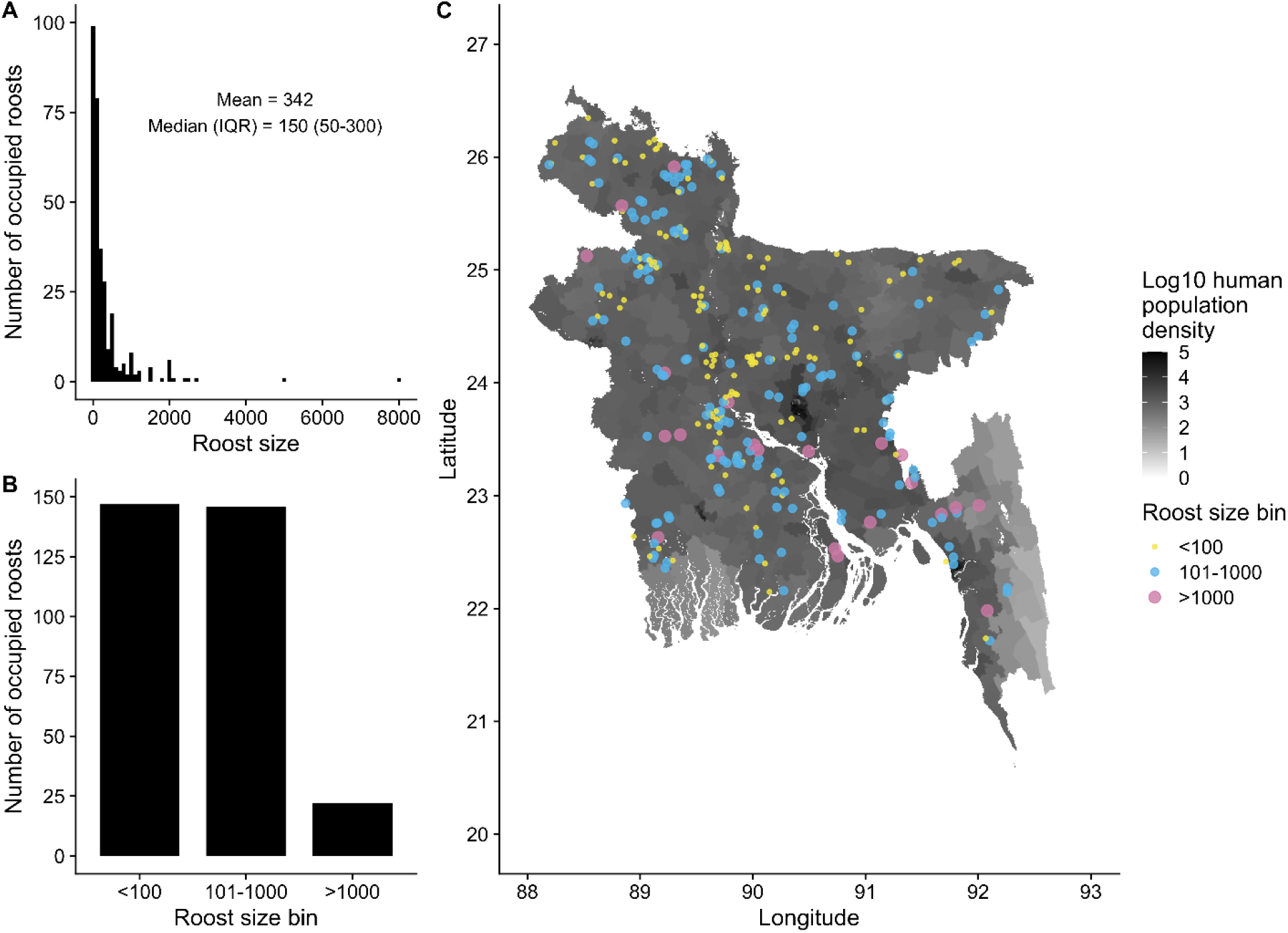
Size and geographic distribution of *Pteropus medius* populations at occupied roost sites (N = 307) in Bangladesh. Roost sizes varied widely from 0 to 8,000 bats (A) but most roosts contained fewer than 1,000 bats (B). Roosts of varying size were observed throughout the country (C) where human population density is high (1,134 persons/km^2^ in the whole country in 2010).

Roost sizes did not appear to be spatially clustered, such that large and small roosts are intermixed throughout the country (Figure 3C). The clustering of roosts in the central and northwest regions of Bangladesh appears to be a spatial artefact of the sampling design, which targeted roost sites predominantly in and nearby villages where Nipah virus spillover events have occurred (Figure A7). Following model selection using lasso, the remaining spatial covariates generally had poor explanatory power for roost occupancy (presence/absence of bats) and abundance (roost size), with R^2^ of 15% or less for testing and training sets (Table 1). AUC was 70% or less for models of occupancy, which indicates poor discriminatory power for predicting occupied and unoccupied roosts [111].

**Table 1.** Performance metrics of GLM and random forests of bat roost occupancy and abundance.

| Response variable | Set | Model | Response error | RMSE | MAE | $R^2$ | AUC |
| --- | --- | --- | --- | --- | --- | --- | --- |
| Occupancy (presence/absence of bats) | Training (n = 380) | GLM | 0.48 | 0.45 | 0.42 | 0.12 | 0.7 |
|  |  | Random forest |  | 0.48 | 0.41 | 0.04 | 0.61 |
|  | Test (n = 94) | GLM | 0.46 | 0.46 | 0.43 | 0.02 | 0.59 |
|  |  | Random forest |  | 0.51 | 0.43 | 0 | 0.49 |
| Abundance (roost size) | Training (n = 255) | GLM | 670 | 631 | 314 | 0.14 |  |
|  |  | Random forest |  | 643 | 312 | 0.09 |  |
|  | Test (n = 60) | GLM | 744 | 711 | 320 | 0.1 |  |
|  |  | Random forest |  | 709 | 327 | 0.08 |  |

RMSE – root mean square error, MAE – mean absolute error, AUC – error under the receiver operating characteristic curve.

These results broadly indicate that bat roosts are not linearly associated with the available covariate data and largely reflect the geography of nearby villages that were surveyed (Tables A5–A6). For example, an average roost site is situated in an area with high human population density, close to inland water bodies, with a nearby feeding site (fruit trees) or date palm tree within 5 km, and numerous feeding sites or date palm trees within a 15 km radius around the site (Table 2; Figure A8). This pattern is consistent with Bangladesh as a whole, where human population density is high everywhere (Figure 3C) and villages contain numerous potential fruit and date palm trees that could attract bats (Figure A7). Only seven out of 474 roost sites had no date palm trees or feeding sites within 15 km of the roost site. However, all of these roost sites had a date palm tree or feeding site within 25 km of the roost site. Roost sizes showed similarly static distributions compared to the other 28 covariates assessed (Table A1; Figures A9–A11). Similar to other studies of *P. medius*, roost sites were close to water bodies (Table 1) [101,102,105], but distance to water did not explain variation in the occupancy or abundance of bats at roost sites (Tables A5–A6).

**Table 2.** Distribution of select spatial covariates across all mapped roost sites.

| Covariate | Median (IQR) |
| --- | --- |
| Human population density (persons/km<sup>2</sup>) | 996 (858–1,260) |
| Distance to nearest inland water (km) | 0.6 (0.3–1) |
| Distance to nearest feeding site (km) | 2 (0.9–3.6) |
| Distance to nearest date palm tree (km) | 1.2 (0.2–2.7) |
| Number of feeding sites within 15 km of roost site | 11 (3–20) |
| Number of date palm trees within 15 km of roost site | 80 (29–307) |

Despite the widespread distribution of bat roost sites and the presence of some relatively large roosts (>1,000 bats), interviewees report that, with respect to their own memory, most roosts are decreasing in size (Figure 4A). These patterns support anecdotal reports of decreasing *P. medius* populations from biologists and bat hunters, a trend attributed to cutting of roost trees and overhunting [66,67]. Local Nipah virus spillover investigation teams have reported that village residents will often cut down roost trees within villages after an outbreak [44]. In support of this, we observed that roost sites in and around Nipah virus case villages had more unoccupied roosts than control villages that were either near (>5 km) or far (>50 km) from case villages (Figure 4B). Besides cutting down roost trees, interviewees listed a number of other reasons that bats left a roost site, including that bats were hunted, or bats were harassed with rocks, mud, sticks, or gunfire (Figure 4C).

**Figure 4.**
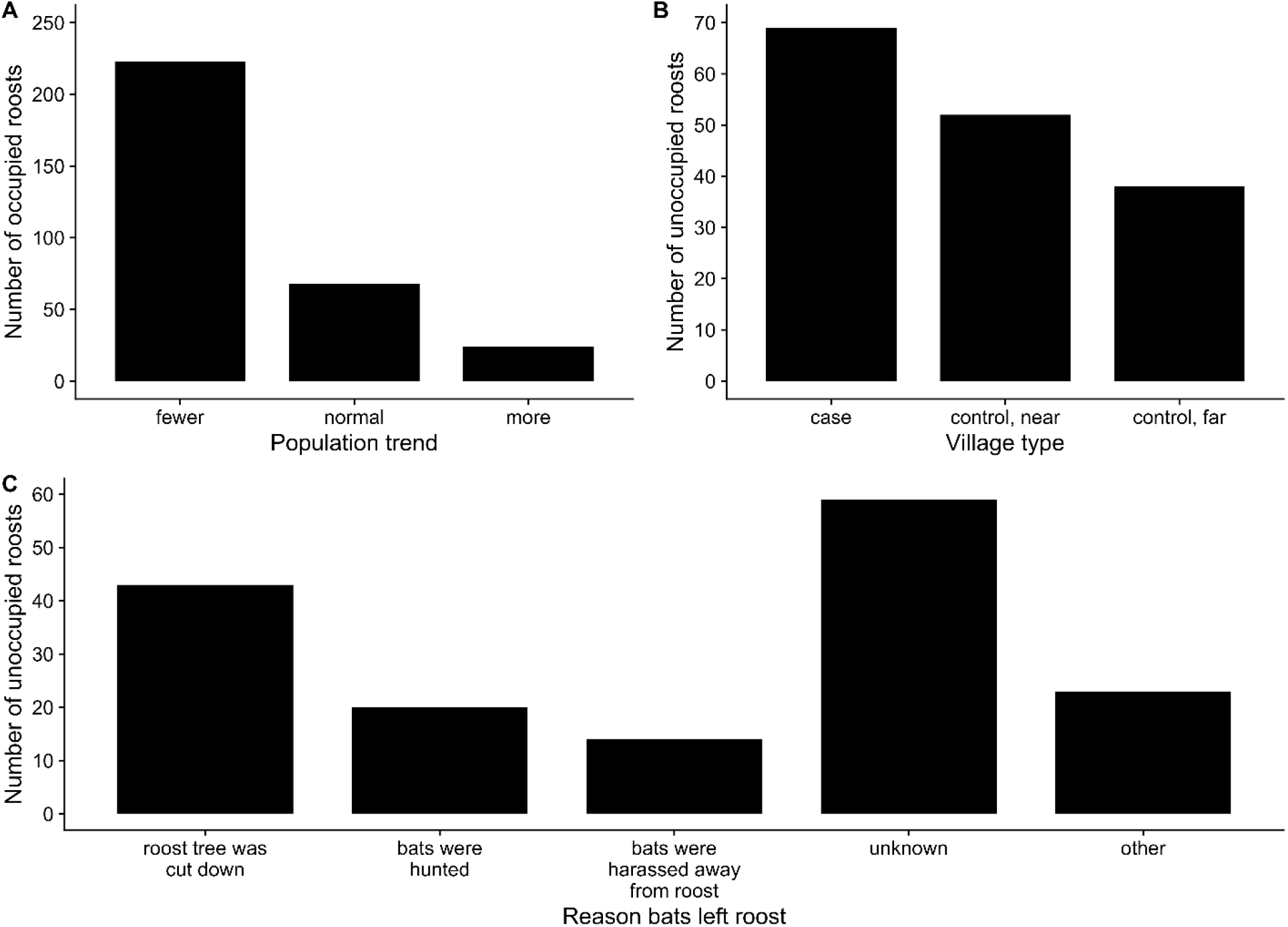
Reported trends for *Pteropus medius* populations at occupied roost sites (A); distribution of unoccupied roost sites across Nipah virus case villages and control villages (B); and reported reasons for bats no longer occupying roost sites (C).

### Historical land use change in Bangladesh

According to the collated data, the majority of forest loss in Bangladesh occurred prior to the 20^th^ century but has steadily continued to the present (Figure 5). Prior to human occupation of the land area comprising Bangladesh, the whole country was likely covered in dense tropical forest, similar to neighboring countries in Southeast Asia [83]. Evidence of human occupation in Bangladesh dates back at least 20,000 years, rice cultivation and domesticated animals occurred before 1500 BCE, and sedentary urban centers were seen by the fifth century BCE [112]. Clearing of land for rice cultivation continued through to the 16^th^ century CE, by which time rice was being exported from the Bengal delta to areas of South and Southeast Asia. During Mughal rule over the Bengal delta starting in the 1610, the Ganges (Padma) River shifted eastward, so Mughal officials encouraged colonists to clear forests and cultivate rice in eastern Bangladesh [112]. Thus, much of the native forests in Bangladesh had been converted to cultivated land prior to 1700 (Figure 5).

**Figure 5.**
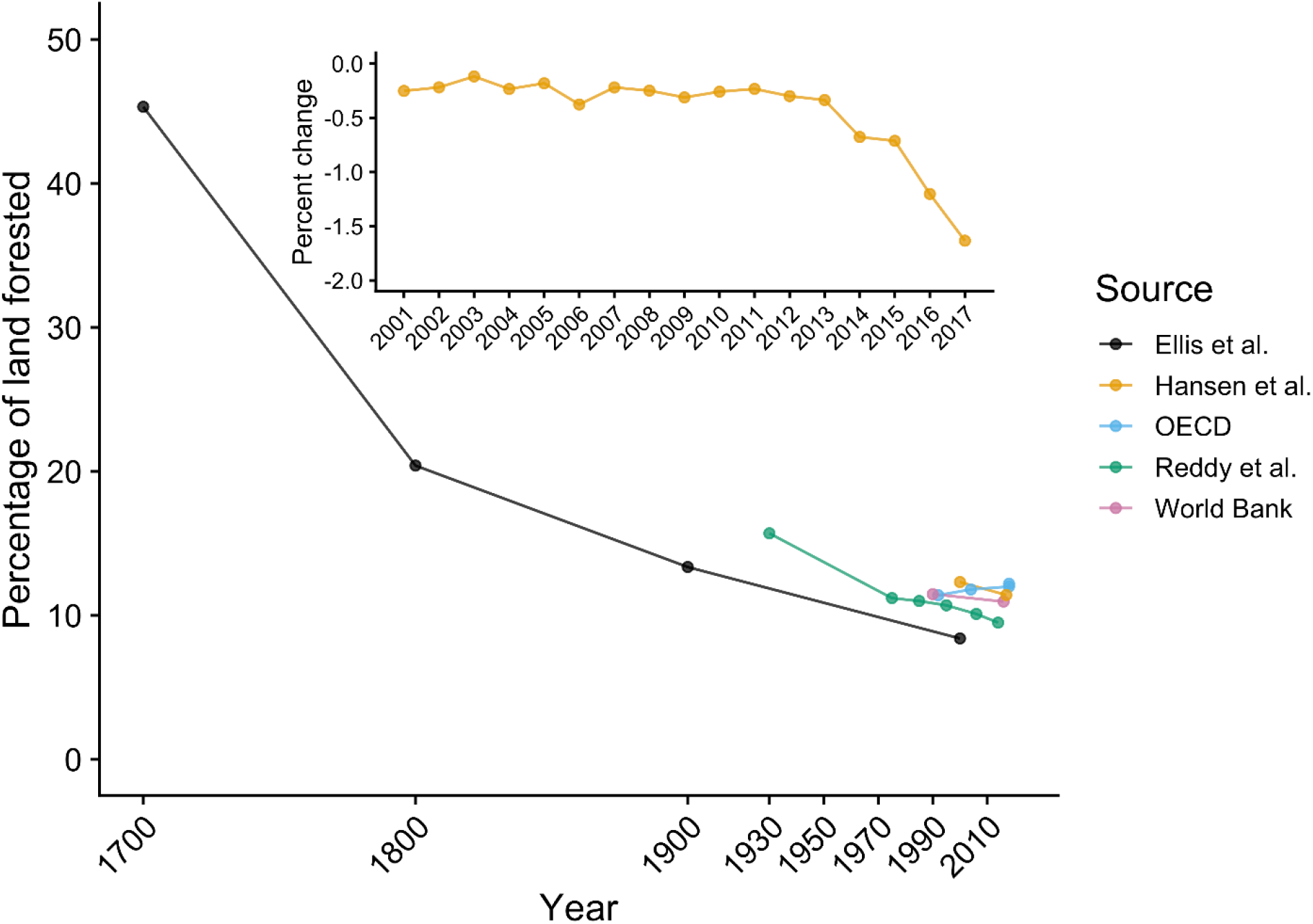
Historical change in forested land area in Bangladesh according to available sources. Inset displays the rate of dense forest loss (annual percent change) since 2000, with a recent increase in this rate of decline, drawn from Hansen et al. [82]. A cutoff value of 40% was used to define dense forests. Only gross forest loss is displayed, since data on forest gain only covers the period 2000–2012.

Following the Battle of Plassey in 1757, the British East India Company took control of the country and established Permanent Settlement, a system of land taxation that set a fixed tax burden for landholders (zamindars). While the intention was that the fixed tax rates would allow zamindars to invest more in agricultural development of the land through better seeds, irrigation, and tools, this never materialized. Since the British would auction the zamindar’s land if they fell behind on their tax obligation, land became a valuable commodity that was bought and sold by wealthy bureaucrats and zamindars. This fostered a hierarchical system where the peasantry working the land paid rent but had no property rights, while landowners were only attached to the land through a series of intermediary managers. To meet their tax obligation and collect rent from tenant farmers, landowners encouraged cultivation of cash crops (cotton, indigo, sugarcane, silk, tea, tobacco, and jute) meant for export in the global market. Agrarian production increased not through agricultural intensification of already cultivated land, but through clearing of native forest. Forest cover declined dramatically during the 1700s and 1800s (Figure 5; Figure A12) and the system of Permanent Settlement existed with some modifications until the 1950s [112].

Production of sugar for export and local consumption came predominantly from sugarcane during the colonial period, but a minor proportion (perhaps 10–15%) was produced from date palm sap from cultivated *Phoenix sylvestris*. While historically date palm sugar was used locally for the preparation of sweetened foods, it became integrated into the global sugar trade starting in 1813 and the value of date palm sap increased. The number of date palms in Bangladesh increased rapidly from the 1830s and remained high until at least the early 1900s, propelled by British encouragement of landowners and the development of mills by the British to produce sugar from date palm sap [61]. Roughly 1,370 metric tons of raw sugar (*gur*) was produced from date palm sap on average during 1792–1813 in Bangladesh, which increased to 38,000 tons of *gur* in 1848 and 162,858 tons by 1905, and then decreased to 66,930 tons by 1911 [61]. The most recent figures from the Bangladesh Bureau of Statistics for 2016–2017 put the area of Bangladesh under date palm cultivation for sap at 20.8 km^2^ with a production of 169,056 metric tons of palm sap (perhaps 10% of which might be converted to *gur*) [113,114]. This is compared to 920 km^2^ under sugarcane producing 3,862,775 tons of sugarcane juice during the same year [113].

Today, Bangladesh has less than 14% of its forest remaining (Figure 5) and the only dense forests are restricted to the southwestern mangrove forests of the Sundarbans and the southeastern forests of the Chittagong Hill Tracts (Figure A12). The portion of the Sundarbans in Bangladesh is a protected as the Sundarban Reserve Forest containing three large wildlife sanctuaries. The region of the Chittagong Hills had enjoyed a level of political autonomy during Mughal rule and was also the last part of Bangladesh to come under state rule after the British invaded in 1860 but retained some regional autonomy in their system of taxation and land rights [112]. Combined with the more rugged terrain of this region, intensification of industrial forestry and agricultural production was delayed until the 1900s, and this region remains one of the least populated areas of the country (Figure 3). These conditions have thereby preserved much of the primary forest until the present (Figure A12). The conditions in neighboring Myanmar were similar, as the British did not begin their rule of the country until 1824. Prior to British rule, Myanmar’s agricultural economy was not as export-focused compared to Bangladesh, but this shifted towards intensified production of rice for export during the colonial period [115]. Partly due to a delayed agricultural intensification imposed by the British, trees still cover around half of Myanmar’s land area [85] and the population density was only 77 persons/km^2^ in 2010 [72].

Recent deforestation in Bangladesh has continued at a steady pace, with a net rate of 0.75% or less per year during 1930–2014 [88], and is concentrated in eastern Chittagong Division (Figure A13). However, there has been a rise in deforestation since 2013 (Figure 5 inset). Additionally, felling of tall trees continued even in largely deforested areas of Bangladesh for the purpose of curing tobacco leaves and brick burning [67]. Since *P. medius* relies on tall tree species such as banyan (*Ficus benghalensis*) to form large roosts [73], the loss of single tall trees can scatter bats into ever smaller populations.

## Discussion

### Historical land use change, bat ecology, and Nipah virus spillover

Given the nearly two decades of research on Nipah virus in Bangladesh, there are facets of its ecology that are now clear. Historical patterns of forest loss have drastically diminished native habitat for fruit bats. *Pteropus medius* bats now live in mostly small, resident roosts in close proximity to humans and opportunistically feed on cultivated food resources. These gradual but dramatic changes have produced a system that facilitates spillover of a bat-borne virus. The consequence is almost annual spillover of Nipah virus in winter months following consumption of raw or fermented date palm sap that has been contaminated with bat excreta containing Nipah virus.

Our analysis suggests that the current state of the bat-human ecological system in Bangladesh supports Nipah virus spillover: a mobile metapopulation of reservoir hosts living amongst humans and sharing food resources that has likely existed for many years prior to the first recognized outbreaks. While the loss of forests in Bangladesh is still occurring and potentially affecting the distribution of *P. medius*, the majority of the land use change from forest to cultivated areas occurred at least a century ago (Figure 5). Cultivation of date palm trees for their sap and other products is a tradition that has likely been practiced for centuries [116], and bats have been potentially consuming sap for an equal amount of time. Importantly, the date palm sap industry was greatly expanded by the British during the late 19^th^ and early 20^th^ centuries and continues at a similar scale to the present [61,113]. Time-calibrated phylogenetic analyses indicate that Nipah virus has been circulating in *P. medius* in Bangladesh and India since the 1950s or earlier [6,117,118]. Thus, none of the conditions that promote Nipah virus spillover in Bangladesh are new. Spillovers almost certainly occurred in the past but were undetected prior to the first isolation of Nipah virus in 1999 and the subsequent development of diagnostic tests. Even recent outbreaks since surveillance was established in 2007 might be missed. Hegde et al. found that because encephalitis case patients are less likely to attend a surveillance hospital if it is distant from their home and if their symptoms are less severe, at least half of all Nipah virus outbreaks during 2007–2014 were likely missed [119].

The ecological state of Nipah virus in Bangladesh has important similarities and differences with the ecology of the related Hendra virus in *Pteropus* spp. in Australia. Spillover events from bats primarily occur in the cooler, dry winter months in both Australia and Bangladesh, and evidence from Australia suggests that this season is when bats are potentially experiencing nutritional stress, are residing in small roosts close to humans, and are shedding more viruses [24,120]. In contrast to *P. medius* in Bangladesh, *Pteropus* populations in Australia exhibit a range of population sizes and behaviors, from large, nomadic groups that track seasonally available nectar sources to small, resident colonies that feed on anthropogenic resources [108]. The increasing incidence of Hendra virus spillovers is linked with periods of acute food shortage that shift bats from nomadism to residency and drive bats to feed on suboptimal food sources, thereby exacerbating stress and associated viral shedding (Eby et al., in review) [121].

We propose that the systems of Nipah virus in Bangladesh and Hendra virus in Australia represent distinct points on a continuum describing patterns of bat aggregation and feeding behavior in a landscape of available roosting sites and food resources (Figure 6). One end of the spectrum is characterized by seasonal shifts from smaller populations to large aggregations of bats in response to transient pulses in fruit and nectar resources (fission-fusion). The other end of the spectrum represents a permanent state of fission, where bats are distributed in small, mostly resident roosts in a matrix of anthropogenic food resources. Bangladesh appears to fall at the latter end of the spectrum, wherein historical land use change and urbanization removed the native forest habitats that supported *Pteropus medius* populations, leaving limited roosting sites but abundant cultivated fruits that are sufficient for sustaining small populations of bats. Australia would traditionally have been on the opposite end of the spectrum, but loss of winter habitat and urban encroachment may be pushing the system towards more permanent fission, which could result in more consistent spillovers of Hendra virus (Eby et al. in review) [121]. Similar anthropogenic pressures acting on pteropodid bat populations in Southeast Asia or Africa could push these systems into a state similar to Bangladesh, consequently increasing the risk of henipavirus spillover [24].

**Figure 6.**
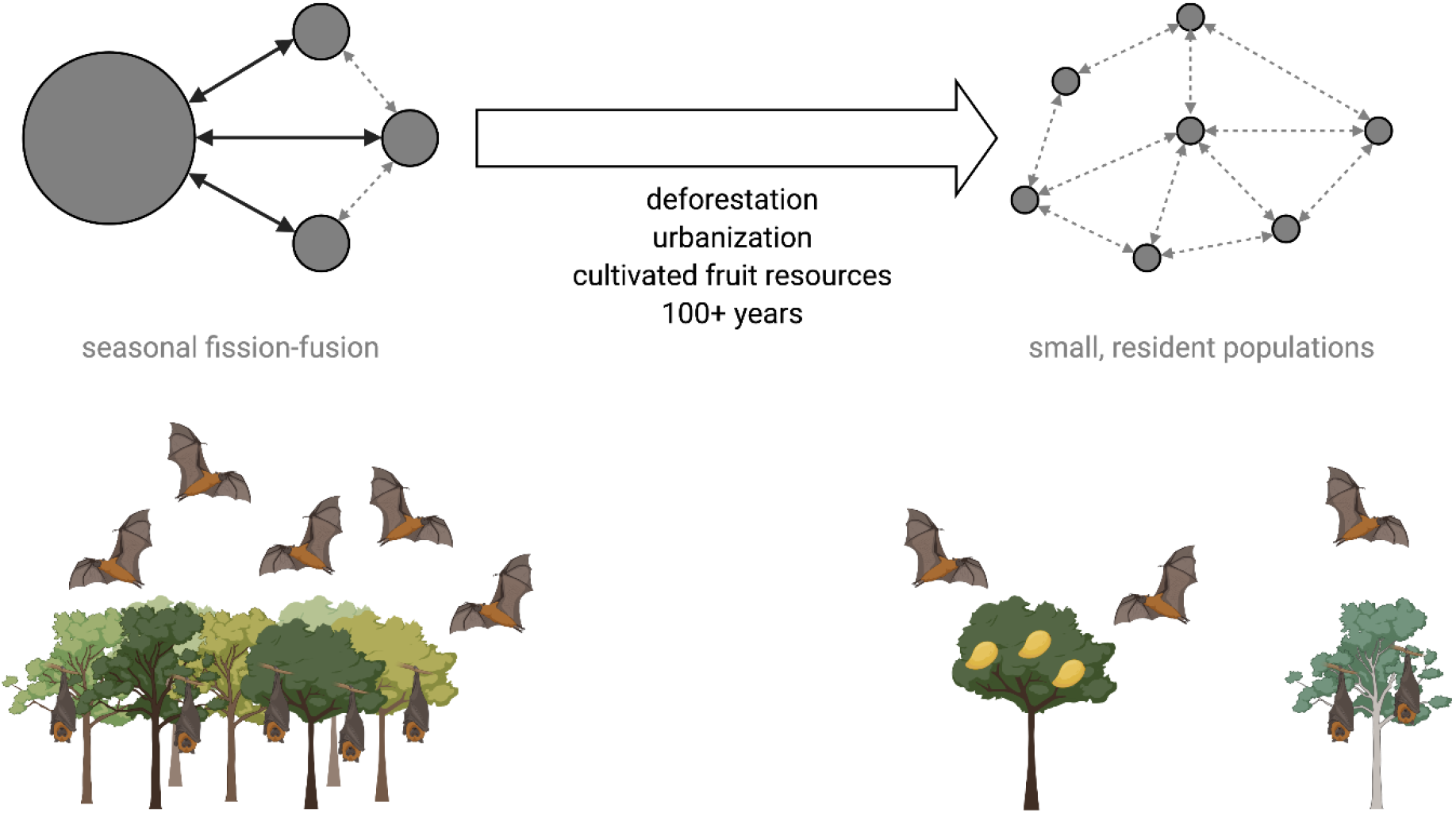
Long-term shifts in pteropodid bat populations and seasonal movements due to anthropogenic land use change. Black arrows show seasonal movements of bats into large aggregations. Dashed gray arrows represent occasional bat movement between roost sites.

### Seasonality of date palm sap consumption and spillovers

Beyond the broad ecological forces that facilitate henipavirus spillover from bats, there are epidemiological patterns that will require further research to explain. Perhaps the most complex are the causes of winter seasonality in Nipah virus spillovers. Recent evidence suggests that bats shed Nipah virus at low levels throughout the year [63]. Date palm trees are also tapped year-round for *tari* production but harvesting increases during winter months to meet increased demand for *tari* and fresh sap [41,43]. Visits by *P. medius* to date palm trees are more frequent in winter [56], even when date palms are tapped year-round for *tari* production (Islam et al., in preparation). Therefore, the risk of viral spillover is always present, but may increase during winter because bats are capitalizing on a resource when it is most available, thereby increasing the probability that sap is contaminated during the winter harvest.

The observation that more Nipah virus spillovers occur during years with colder winters indicates that climate is affecting one or more factors in the system: date palm physiology, bat and human behavior, bat physiology and immunology that affect viral replication, or some combination of these factors. Date palm sap collectors report that date palm sap is sweeter and flows more freely during cooler weather [43,56,61]. These might be physiological responses of *Phoenix sylvestris* to seasonal weather conditions (e.g., sugar or water is concentrated in the trunk during cool, dry weather), yet no data are available on variation in sap flow or sugar content for this species outside of winter months [61]. Harvesting date palm sap when it is sweetest would be optimal not only for the collectors, but also for bats. Fewer cultivated fruits are available during winter than other seasons [58], so bats may gravitate towards date palms because it is readily available during a time of relative food scarcity. More surveys of *P. medius* feeding behavior and the fruits they consume at different times of the year would be necessary to assess this hypothesis [122]. Complementary experiments could be performed to evaluate whether pteropodid bats perceive small differences in sugar concentration and modify their feeding behavior in response to varying energy demands [123].

Another hypothesis, derived from research on Hendra virus in Australian bats, posits that bats shed viruses more frequently during periods of nutritional stress that compromise bat immune function [24,124]. Increased metabolic demands of thermoregulation during winter when food resources are already limited could produce physiological and nutritional stress in bats. Bats may seek out alternative foods (e.g., date palm sap) to compensate for this stress. Whether *P. medius* are shedding more Nipah virus when they are experiencing physiological or nutritional stress in winter is an open question. We need more documentation of body condition, biomarkers of stress and immune function, or abortion rates among female bats to understand any relationships between Nipah virus shedding, stress, and climate [24,125–127].

We also lack information on how seasonal bat movements might influence Nipah virus spillover dynamics. Although our data suggest that most roost sites are continuously occupied (Figure 2), there may still be some seasonal dynamics in bat population sizes as individuals make occasional movements to use seasonally available resources or aggregate for mating. There is evidence from India and Nepal that *P. medius* roost populations vary seasonally, with larger populations in fall and winter than in summer [128,129]. This is mirrored by our data showing winter is the season when more roosts were founded, and bats are present at intermittently occupied sites (Figure A6). There is also evidence that *P. medius* home ranges contract during the dry season (including winter) than the wet season [63]. Nevertheless, genetic data on *P. medius* and Nipah virus in Bangladesh indicate that bat movements are common enough to promote genetic admixture and spread distinct Nipah virus genotypes among geographically distant *P. medius* populations [6]. To better understand how bat movements influence spillover dynamics, we need more information on seasonal variation in bat population sizes at roost sites and potentially individual movement tracking data, which could be used to parameterize metapopulation models of Nipah virus transmission.

### *Roost tree loss and* Pteropus *roosting behavior*

In addition to the causes of seasonality in Nipah virus spillover, more research is needed to determine the effects of current deforestation and human disturbance on *P. medius* populations. While historical patterns of deforestation and land use change have undoubtedly reduced available habitat for pteropodid bats (Figure 5), the effects of current deforestation may be easiest to measure at the scale of individual roost trees. If a single tree in a largely deforested area has qualities that are preferred by bats and therefore supports a large population of bats, loss of that tree could have a very large effect on the bat population but would contribute very little to overall deforestation rates. Our statistical analysis was unable to explain substantial variation in the occupancy and size of roosts using available data on spatial covariates, including land use, human population density, bioclimatic variables, and distribution of cultivated fruit and date palm trees (Table 1; Table A1). Similar results were observed for *P. medius* populations in Uttar Pradesh, India [101]. Kumar and Elangovan [101] were unable to explain variation in colony size using data on distance to human settlements, roads, or water bodies. However, they did find that colony size increased with tree height, trunk diameter, and canopy spread. The majority of colonies were found in tree species with wide canopies, including *Ficus* spp., mango, *Syzygium cumini*, and *Madhuca longifolia* [101]. Hahn et al. [73] compared occupied roost trees to non-roost trees within a 20×20 m area around central roost trees and found that *P. medius* in Bangladesh favor tall canopy trees with large trunk diameters. Therefore, future efforts to understand variation in *P. medius* population sizes across Bangladesh should collect more data on characteristics of roost trees. Furthermore, the sampling design of our population meant that no bat roosts could have been observed further than 5 km from a village, meaning that bat roosts in remnant forested areas in the Sundarbans and Chittagong Hills were much less likely to be included in the study (Figure A7). Further surveys of roost sites may reveal distinct roosting patterns of *P. medius* populations living in these areas or in other areas within the range of *P. medius* where human population density is lower and forested habitat is more intact.

Our survey data also indicate that many roost sites are frequently abandoned following harassment, hunting, or removal of roost trees and that more unoccupied roosts are found near villages that have experienced Nipah virus spillover (Figure 4). Presumably these bats disperse and form new roosts or join existing roosts, but the new roost trees may be of lower quality than the previous roost and only support a smaller population of bats. More granular data on the cumulative effects of roost tree loss on average *P. medius* population sizes would refine our conceptual model of shifting roosting behavior in pteropodid bats (Figure 6). Moreover, movements of bats following abandonment of roost sites could have implications for Nipah virus transmission dynamics. Dispersal of bats following roost tree loss or harassment could lead infected bats to seed outbreaks elsewhere [124]. Therefore, reactionary cutting of roost trees in villages with Nipah virus spillovers is counterproductive for spillover prevention and bat conservation and should be discouraged.

### Possible interventions to prevent Nipah virus spillover

Finally, there is a need to explore possible interventions to prevent Nipah virus spillover. Without a vaccine for Nipah virus, much of the research has focused on mitigating the risk of spillovers. Several studies in Bangladesh have centered on educating the public about the risks of drinking raw date palm sap and methods for preventing bat access to date palm sap during collection [130–132]. There is also a need for increased surveillance of bats and humans in close contact with bats in Bangladesh and other areas within the range of *Pteropus* bats. These enhanced surveillance efforts could include serosurveys of bat hunters, date palm sap collectors, people who drink sap or eat fruits that have been partially consumed by bats, and people who live in close proximity to bat roost sites [13,66,133,134]. While there has been no evidence that consuming fruits partially eaten by bats is associated with Nipah virus spillover to humans in Bangladesh and Cambodia [13,135], this route was believed to be the cause of the 1998–1999 outbreaks in pigs that led to human cases in Malaysia and Singapore [54]. A 2009 survey of livestock in Bangladesh living nearby to *Pteropus* bat roosts also found henipavirus antibodies in 6.5% of cattle, 4.3% of goats, and 44.2% of pigs [136]. Animals were more likely to be seropositive if they had a history of feeding on fruits partially eaten by bats or birds and drinking date palm juice from Asian palmyra palms (*Borassus flabellifer*) [136]. Therefore, Nipah virus transmission from livestock to humans in Bangladesh is a risk that should be explored with additional serosurveys and efforts to limit contact of livestock with fruits and other materials potentially contaminated with bat excreta.

Similar risks may apply in neighboring India where Nipah virus outbreaks have been linked to fruit bats [48,137]. The index case of a 2007 Nipah outbreak in West Bengal was reported to frequently drink date palm liquor (*tari*) and had numerous bats living in trees around their home [48]. Researchers speculate that the 2018 and 2019 outbreaks in Kerala, India, may be linked to consumption of partially eaten fruits [137]. However, this has not been confirmed via detection of Nipah virus on partially eaten fruits or case-control studies [39,44]. The index case associated with 23 cases of Nipah virus infection during the 2018 Kerala outbreak reported possible contact with an infected baby bat, but this was also not confirmed [39]. Silver date palm is not cultivated for sap in Kerala, but coconut palm and Asian palmyra palm are [39]. The narrow-mouthed containers that are used to collect sap from these palm species are thought to prevent bat access to the sap within the container [39] but might not prevent bats from accessing and contaminating sap at the tapping site or from inflorescences. Additional studies using infrared cameras to understand fruit bat feeding behavior around other palm trees harvested for sap and possible intervention methods similar to those done in Bangladesh are warranted [56,130]. Such information would help to clarify how Nipah virus is transmitted from fruit bats to humans in India and allow for ecological comparison of outbreaks in these two neighboring countries.

At a higher level, methods that limit human-bat contact through ecological interventions may be beneficial. Plantations of fruit- and nectar-producing tree species could provide alternative food for *P. medius*, such as cotton silk (*Ceiba petandra, Bombax ceiba*), Indian mast tree (*Polyalthia longifolia*), and Singapore cherry (*Muntingia calabura*). Trees that produce fruit year-round or specifically during winter could provide bats with the required nutrition that would have been acquired from date palm sap or other cultivated fruits. In combination with methods to prevent bat access to date palm sap, ecological interventions that would allow *P. medius* populations to persist in Bangladesh and other areas while lowering the risk of Nipah virus spillover should be explored.

## Conclusions

The ecological conditions that produce yearly spillovers of Nipah virus in Bangladesh are not a new phenomenon, but rather a culmination of centuries of anthropogenic change. The opportunistic feeding behavior of *P. medius* has allowed populations to adapt to these modified landscapes, persisting in small, resident colonies feeding on cultivated fruits. Shared use of date palm sap by bats and humans is a key route for Nipah virus spillover during winter months. Continued research on this system could reveal how bat behavior and physiology influence the seasonality of Nipah spillovers and explore potential ecological interventions to prevent spillover.

## Supplementary Materials

The following are available online at www.mdpi.com/xxx, Appendix A: Supplementary tables and figures.

## Author Contributions

Conceptualization, E.G., R.P., and P.H.; data curation, C.M., E.G., and H.S.; formal analysis, C.M.; visualization, C.M.; writing – original draft preparation, C.M.; writing – reviewing and editing, all authors. All authors have read and agreed to the published version of the manuscript.

## Acknowledgments

We thank Peggy Eby and Birgit Nikolay for early discussions on data sources and analyses.

