## Appendix A for "The ecology of Nipah virus in Bangladesh: a nexus of land use change and opportunistic feeding behavior in bats"

**Table A1.** Source information and distribution of spatial covariates across all mapped roost sites.

| Covariate | Source | Timespan | Raster resolution | Median (IQR) |
|---|---|---|---|---|
| BIO1 – annual mean temperature (°C) | WorldClim [1] | 1970–2000 | 1 km | 25.4 (25.1–25.7) |
| BIO2 – mean diurnal temperature range (°C) | | | | 9.7 (9.4–10) |
| BIO5 – maximum temperature of warmest month (°C) | | | | 33.8 (33–34.6) |
| BIO6 – minimum temperature of coldest month (°C) | | | | 11.2 (10.8–11.6) |
| BIO12 – annual precipitation (mm) | | | | 1,937 (1,760–2,281) |
| BIO14 – precipitation of driest month (mm) | | | | 5 (3–8) |
| BIO15 – precipitation seasonality (coefficient of variation) | | | | 92.5 (86.2–98.5) |
| BIO18 – precipitation of warmest quarter (mm) | | | | 952 (689–1,148) |
| Distance to nearest artificial surface (km) | WorldPop/ESA-CCI-LC [2,3] | 2011 | 100 m | 8.1 (4.8–12.5) |
| Distance to nearest bare area (km) | | | | 28.9 (12.9–46.7) |
| Distance to nearest herbaceous area (km) | | | | 11.3 (5.5–20.7) |
| Distance to nearest shrub area (km) | | | | 50.8 (25.3–64.8) |
| Distance to nearest sparse vegetation area (km) | | | | 55.5 (32.7–81) |
| Distance to nearest tree area (km) | | | | 5.1 (2.2–9.3) |
| Distance to nearest inland water (km) | WorldPop/ESA-CCI-LC [2,3] | 2000–2012 | 100 m | 0.6 (0.3–1) |
| Distance to nearest waterway (km) | WorldPop/OSM [2,3] | 2016 | 100 m | 4.5 (1.3–8.5) |
| Distance to nearest road intersection (km) | | | | 4 (2.3–6.7) |
| Distance to nearest road (km) | | | | 1.3 (0.5–2.6) |
| Distance to protected wilderness (km) | WorldPop/IUCN [2,3] | 2000–2017 | 100 m | 197 (148–241) |

| | | | | |
|---|---|---|---|---|
| Human population density (/sq km) | SEDAC/GPW [4] | 2010 | 1 km | 996 (858–1,260) |
| Elevation (m above sea level) | WorldPop [2,3] | 2000 | 100 m | 16 (12–24) |
| Slope | WorldPop [2,3] | 2000 | 100 m | 1 (0–1) |
| Night-time lights (VIIRS) | WorldPop [2,3] | 2012 | 100 m | 0.3 (0.2–0.5) |
| Forest pixels (>10% cover) within 15 km radius | Global Forest Change [5] | 2000 | 30 m | 60,151 (35,557–100,047) |
| Distance to nearest roost site (km) | This study | 2011–2013 | Points | 1.5 (0.2–3.1) |
| Distance to nearest village (km) | | | | 1.8 (0.9–3.2) |
| Distance to nearest feeding site (km) | | | | 2 (0.9–3.6) |
| Distance to nearest date palm tree (km) | | | | 1.2 (0.2–2.7) |
| Roost sites within 15 km radius | | | | 7 (3–13) |
| Villages within 15 km radius | | | | 2 (1–4) |
| Feeding sites within 15 km radius | | | | 11 (3–20) |
| Date palm trees within 15 km radius | | | | 80 (29–307) |

ESA-CCI-LC – European Space Agency Climate Change Initiative land cover, OSM – OpenStreetMap, IUCN – International Union for Conservation of Nature strict nature reserves and wilderness areas, GPW – Gridded Population of the World

**Table A2.** Selection of generalized linear models (GLM) for the number of districts affected by Nipah virus spillover based on selection by AICc. Only models with ΔAICc < 4 are shown.

| **Model** | **DF** | **AICc** | **ΔAICc** |
|---|---|---|---|
| days_below17 | 2 | 85.219 | 0 |
| days_below17 + DMI | 3 | 87.867 | 2.648 |
| days_below17 + MEI | 3 | 88.024 | 2.805 |
| days_below17 + SIOD | 3 | 88.069 | 2.850 |
| temp_mean + days_below17 | 3 | 88.126 | 2.907 |
| temp_min + days_below17 | 3 | 88.127 | 2.908 |
| days_below17 + precip | 3 | 88.133 | 2.914 |

**Table A3.** Selection of generalized linear models (GLM) for the number of Nipah virus spillover events based on selection by AICc. Only models with ΔAICc < 4 are shown.

| Model | DF | AICc | ΔAICc |
|---|---|---|---|
| days_below17 | 2 | 100.331 | 0 |
| days_below17 + precip | 3 | 102.867 | 2.537 |
| days_below17 + DMI | 3 | 102.914 | 2.583 |
| temp_mean + days_below17 | 3 | 102.947 | 2.616 |
| days_below17 + MEI | 3 | 102.991 | 2.661 |
| temp_min + days_below17 | 3 | 103.212 | 2.882 |
| days_below17 + SIOD | 3 | 103.243 | 2.912 |

**Table A4.** Sensitivity analysis for the association between the annual number of Nipah spillovers and the percentage of winter days below a temperature threshold, varying the threshold from 15 to 20 °C. The coefficient is the estimated coefficient for a Poisson GLM. All associations were statistically significant at the 0.01 level (**) or the 0.001 level (***).

| **Outcome** | **Covariate** | **2001–2018** | | **2007–2018** | |
|---|---|---|---|---|---|
| | | **$R^2$** | **Coefficient** | **$R^2$** | **Coefficient** |
| Total_districts | days_below15 | 0.4 | $0.12^{***}$ | 0.82 | $0.13^{***}$ |
| | days_below16 | 0.47 | $0.11^{***}$ | 0.83 | $0.11^{***}$ |
| | days_below17 | 0.53 | $0.11^{***}$ | 0.7 | $0.09^{***}$ |
| | days_below18 | 0.48 | $0.09^{***}$ | 0.6 | $0.07^{***}$ |
| | days_below19 | 0.28 | $0.07^{***}$ | 0.45 | $0.06^{***}$ |
| | days_below20 | 0.13 | $0.05^{**}$ | 0.3 | $0.06^{**}$ |
| Total_events | days_below15 | 0.35 | $0.13^{***}$ | 0.78 | $0.14^{***}$ |
| | days_below16 | 0.43 | $0.12^{***}$ | 0.85 | $0.13^{***}$ |
| | days_below17 | 0.53 | $0.12^{***}$ | 0.79 | $0.11^{***}$ |
| | days_below18 | 0.49 | $0.1^{***}$ | 0.72 | $0.09^{***}$ |
| | days_below19 | 0.29 | $0.07^{***}$ | 0.55 | $0.08^{***}$ |
| | days_below20 | 0.12 | $0.05^{***}$ | 0.33 | $0.07^{***}$ |

**Table A5.** Estimated coefficients for relationships between spatial covariates and bat roost occupancy (presence/absence of bats). Statistical significance of covariates based on estimated coefficients for the test data are shown as: not significant (NS) or significant at the 0.05 level (*), at the 0.01 level (**), or the 0.001 level (***).

| **Covariate** | **Lasso regression coefficient for training data (n = 380)** | **GLM coefficient for test data (n = 94)** | **GLM coefficient t-statistic** |
|---|---|---|---|
| Intercept | 0.69 | 0.95 | $3.8^{***}$ |
| BIO1 – annual mean temperature (°C) | 0 | | |
| BIO2 – mean diurnal temperature range (°C) | 0 | | |
| BIO5 – maximum temperature of warmest month (°C) | -0.028 | -0.76 | $-1.5^{NS}$ |
| BIO6 – minimum temperature of coldest month (°C) | 0 | | |
| BIO12 – annual precipitation (mm) | 0 | | |
| BIO14 – precipitation of driest month (mm) | 0 | | |
| BIO15 – precipitation seasonality (coefficient of variation) | 0 | | |
| BIO18 – precipitation of warmest quarter (mm) | 0.33 | 0.035 | $0.079^{NS}$ |
| Distance to nearest artificial surface (km) | 0 | | |
| Distance to nearest bare area (km) | 0.018 | 0.18 | $0.46^{NS}$ |
| Distance to nearest herbaceous area (km) | 0 | | |
| Distance to nearest shrub area (km) | 0 | | |
| Distance to nearest sparse vegetation area (km) | 0 | | |
| Distance to nearest tree area (km) | 0 | | |
| Distance to nearest inland water (km) | 0.05 | 0.0092 | $0.026^{NS}$ |
| Distance to nearest waterway (km) | -0.17 | 0.2 | $0.65^{NS}$ |

| | | | |
|---|---|---|---|
| Distance to nearest road intersection (km) | 0 | | |
| Distance to nearest road (km) | 0 | | |
| Distance to protected wilderness (km) | 0 | | |
| Human population density (/sq km) | 0 | | |
| Elevation (m above sea level) | 0 | | |
| Slope | 0 | | |
| Night-time lights (VIIRS) | 0 | | |
| Forest pixels (>10% cover) within 15 km radius | 0 | | |
| Distance to nearest roost site (km) | 0 | | |
| Distance to nearest village (km) | 0 | | |
| Distance to nearest feeding site (km) | 0 | | |
| Distance to nearest date palm tree (km) | 0 | | |
| Roost sites within 15 km radius | 0 | | |
| Villages within 15 km radius | 0 | | |
| Feeding sites within 15 km radius | 0 | | |
| Date palm trees within 15 km radius | -0.061 | 0.18 | $0.59^{NS}$ |

**Table A6.** Estimated coefficients for relationships between spatial covariates and bat abundance (roost size). Statistical significance of covariates based on estimated coefficients for the test data are shown as: not significant (NS) or significant at the 0.05 level (*), at the 0.01 level (**), or the 0.001 level (***).

| **Covariate** | **Lasso regression coefficient for training data (n = 255)** | **GLM coefficient for test data (n = 60)** | **GLM coefficient t-statistic** |
|---|---|---|---|
| Intercept | 5.8 | 5.7 | $23.7^{***}$ |
| BIO1 – annual mean temperature (°C) | | | |
| BIO2 – mean diurnal temperature range (°C) | | | |
| BIO5 – maximum temperature of warmest month (°C) | | | |
| BIO6 – minimum temperature of coldest month (°C) | | | |
| BIO12 – annual precipitation (mm) | | | |
| BIO14 – precipitation of driest month (mm) | | | |
| BIO15 – precipitation seasonality (coefficient of variation) | | | |
| BIO18 – precipitation of warmest quarter (mm) | -0.2 | -0.38 | $-1.9^{NS}$ |
| Distance to nearest artificial surface (km) | | | |
| Distance to nearest bare area (km) | | | |
| Distance to nearest herbaceous area (km) | 0.076 | 0.33 | $1.4^{NS}$ |
| Distance to nearest shrub area (km) | | | |
| Distance to nearest sparse vegetation area (km) | | | |
| Distance to nearest tree area (km) | | | |
| Distance to nearest inland water (km) | | | |
| Distance to nearest waterway (km) | 0.072 | 0.31 | $1.3^{NS}$ |

| | | | |
|---|---|---|---|
| Distance to nearest road intersection (km) | | | |
| Distance to nearest road (km) | -0.0046 | 0.05 | $0.19^{NS}$ |
| Distance to protected wilderness (km) | -0.058 | -0.5 | $-1.7^{NS}$ |
| Human population density (/sq km) | | | |
| Elevation (m above sea level) | | | |
| Slope | 0.055 | -0.38 | $-1.4^{NS}$ |
| Night-time lights (VIIRS) | | | |
| Forest pixels (>10% cover) within 15 km radius | 0.19 | 0.33 | $1.7^{NS}$ |
| Distance to nearest roost site (km) | 0.038 | -0.18 | $-0.87^{NS}$ |
| Distance to nearest village (km) | 0.072 | 0.32 | $1.3^{NS}$ |
| Distance to nearest feeding site (km) | | | |
| Distance to nearest date palm tree (km) | | | |
| Roost sites within 15 km radius | -0.11 | -0.11 | $-0.4^{NS}$ |
| Villages within 15 km radius | | | |
| Feeding sites within 15 km radius | | | |
| Date palm trees within 15 km radius | | | |

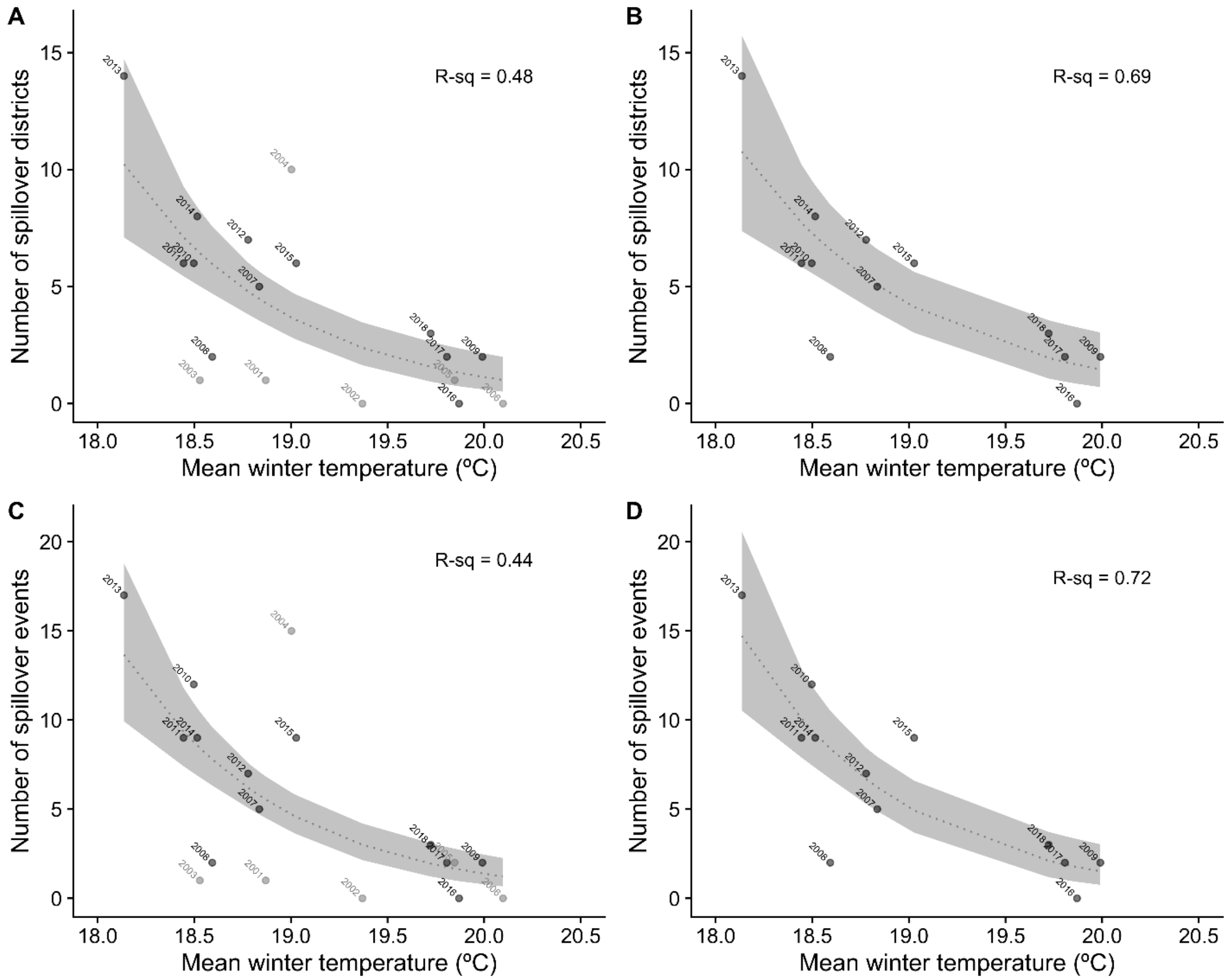


**Figure A1.** Variation in the number of Nipah spillover districts and events explained by mean winter temperatures. Panels show patterns for 2001–2018 (A, C) and 2007–2018 (B, D).

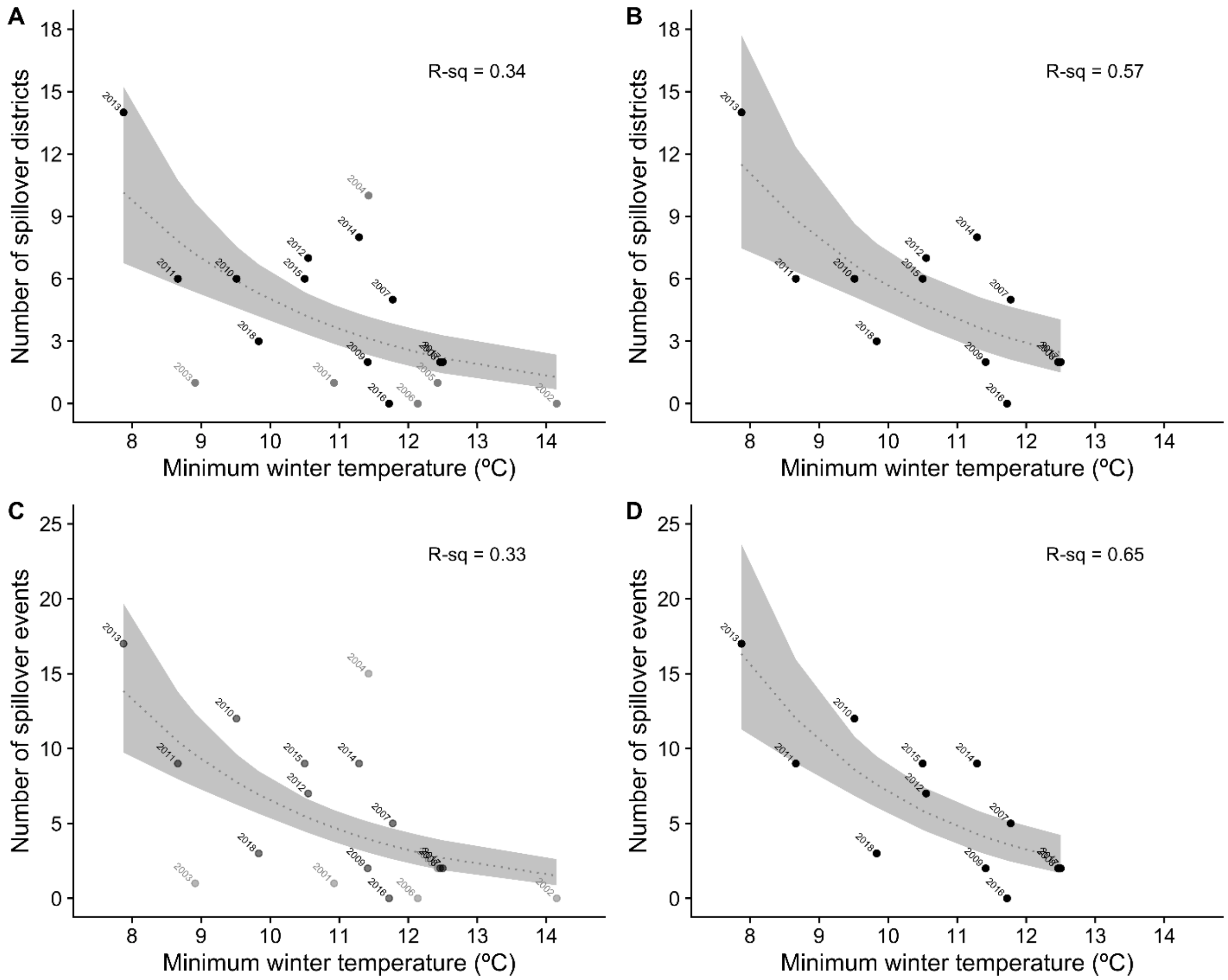


**Figure A2**. Variation in the number of Nipah spillover districts and events explained by minimum winter temperatures. Panels show patterns for 2001–2018 (A, C) and 2007–2018 (B, D).

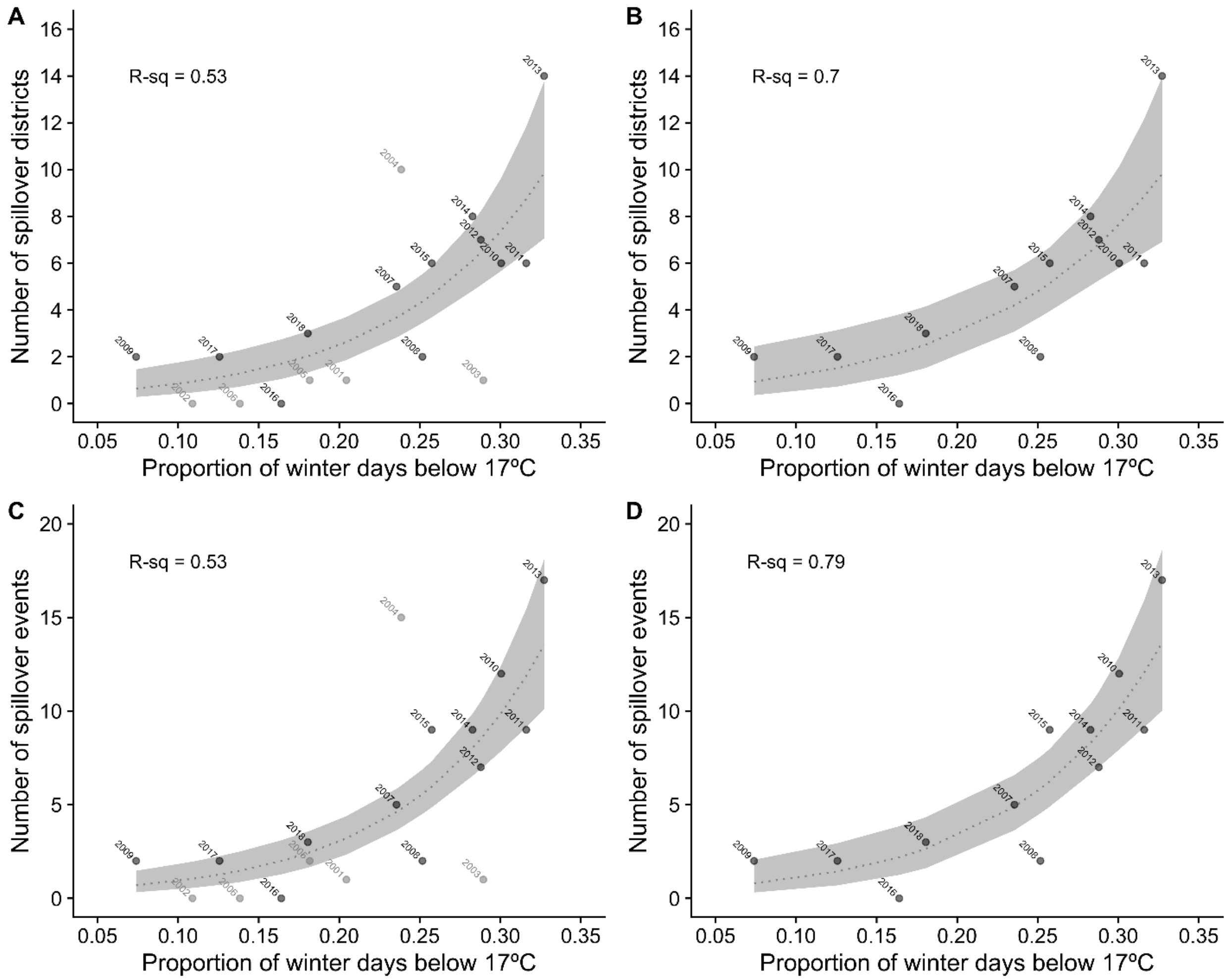


**Figure A3.** Variation in the number of Nipah spillover districts and events explained by cold winter temperatures. Panels show patterns for 2001–2018 (A, C) and 2007–2018 (B, D).

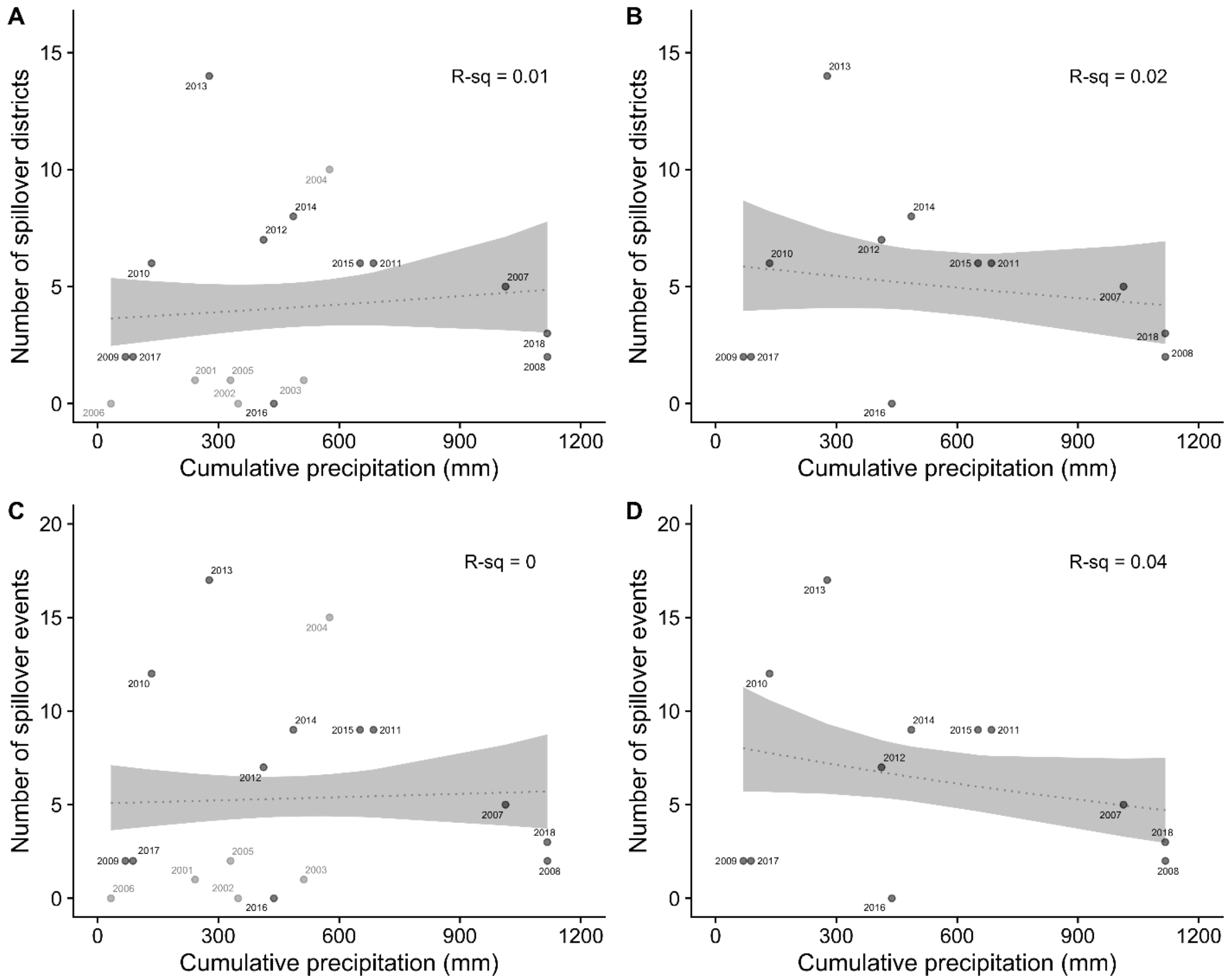


**Figure A4.** Variation in the number of Nipah spillover districts and events explained by cumulative winter precipitation. Panels show patterns for 2001–2018 (A, C) and 2007–2018 (B, D).

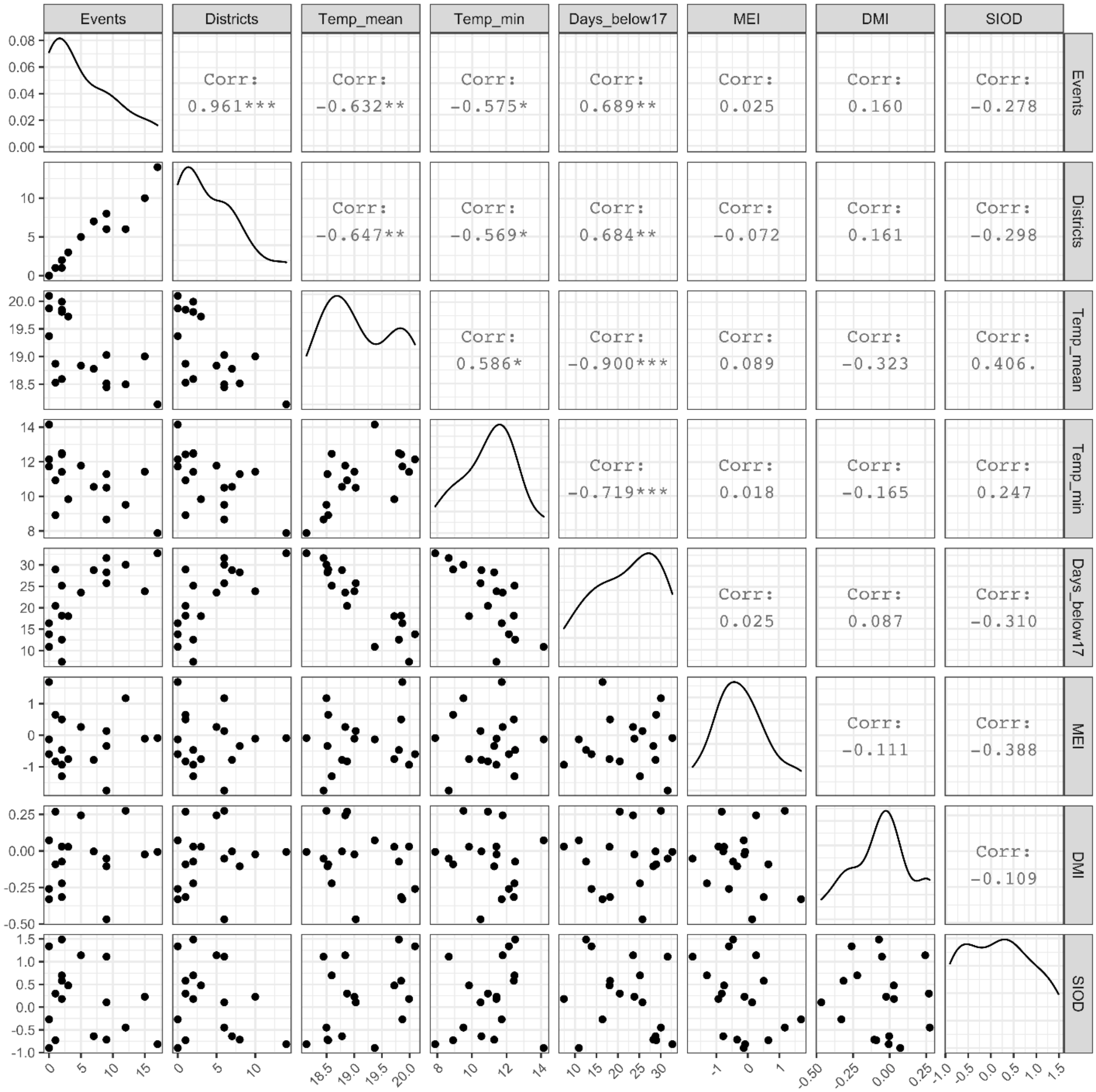


**Figure A5.** Pairwise Pearson's correlation between annual Nipah spillover events, spillover districts, and winter climate measures: mean temperature, minimum temperature, percentage of days below 17 °C, cumulative precipitation, and three indices of climate oscillations (MEI, DMI, and SIOD). Correlations with asterisks are statistically significant at the 0.05 level (*), the 0.01 level (**), and the 0.001 level (***).

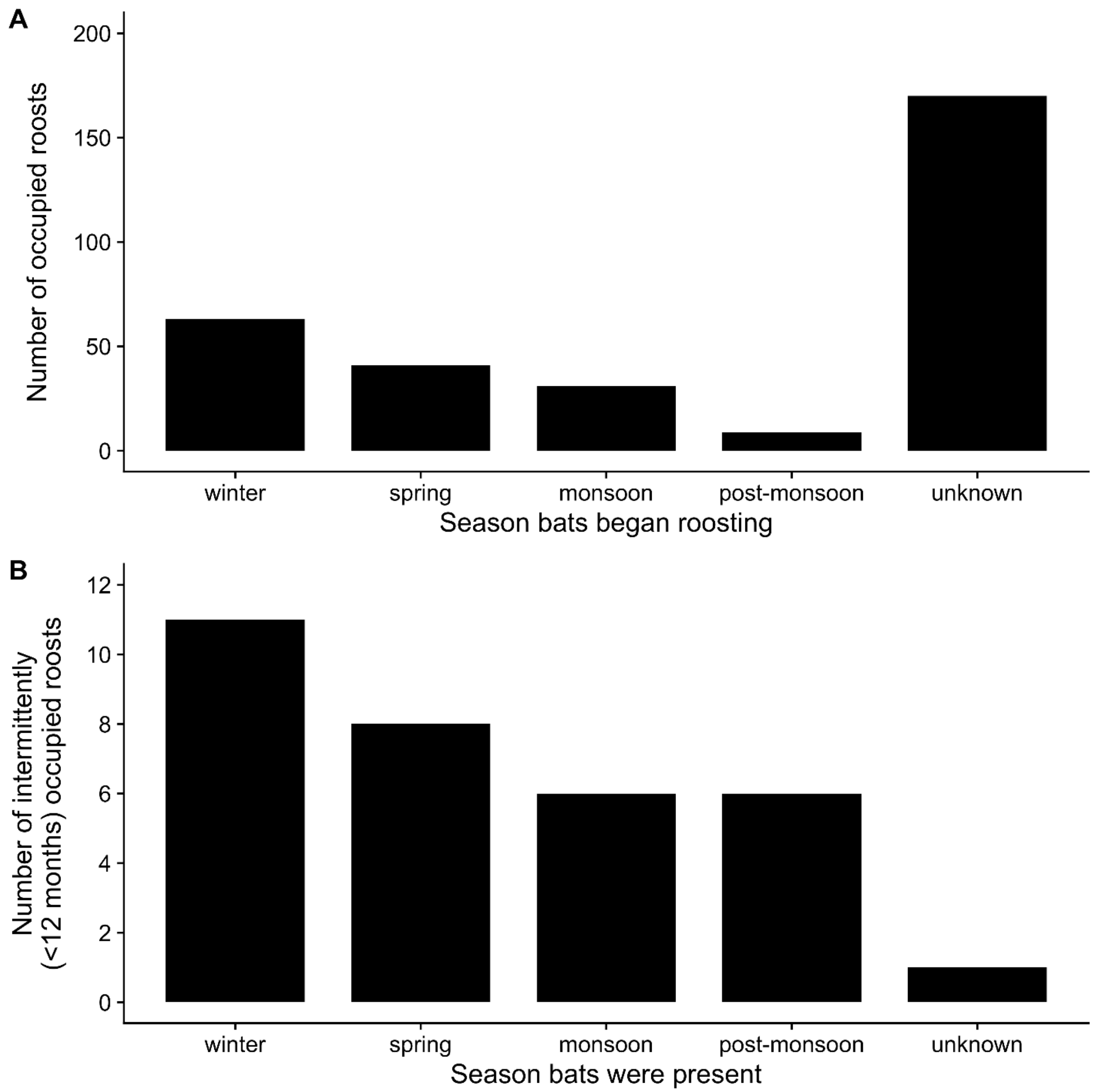


**Figure A6.** Seasonality of *Pteropus medius* roost site occupancy. Panel A shows the reported season when bats began roosting at the site. Panel B shows the season when bats were present at intermittently occupied roost sites (i.e., roosts were occupied <12 months of a year).

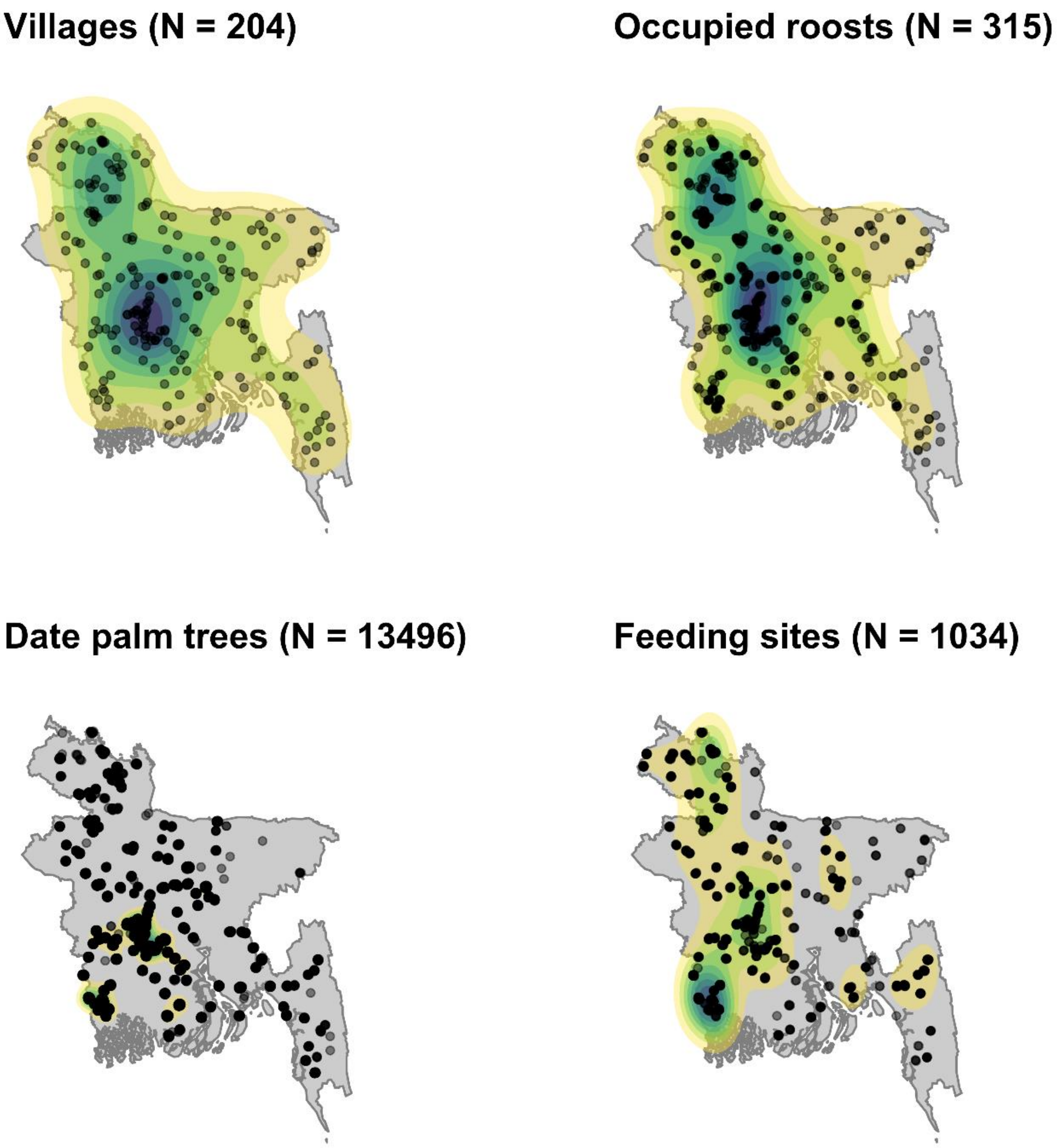


**Figure A7.** Spatial density of study villages, roosts, date palm trees, and bat feeding sites (fruit trees in and around villages). Color contours show the spatial density of events estimated with a bivariate normal kernel.

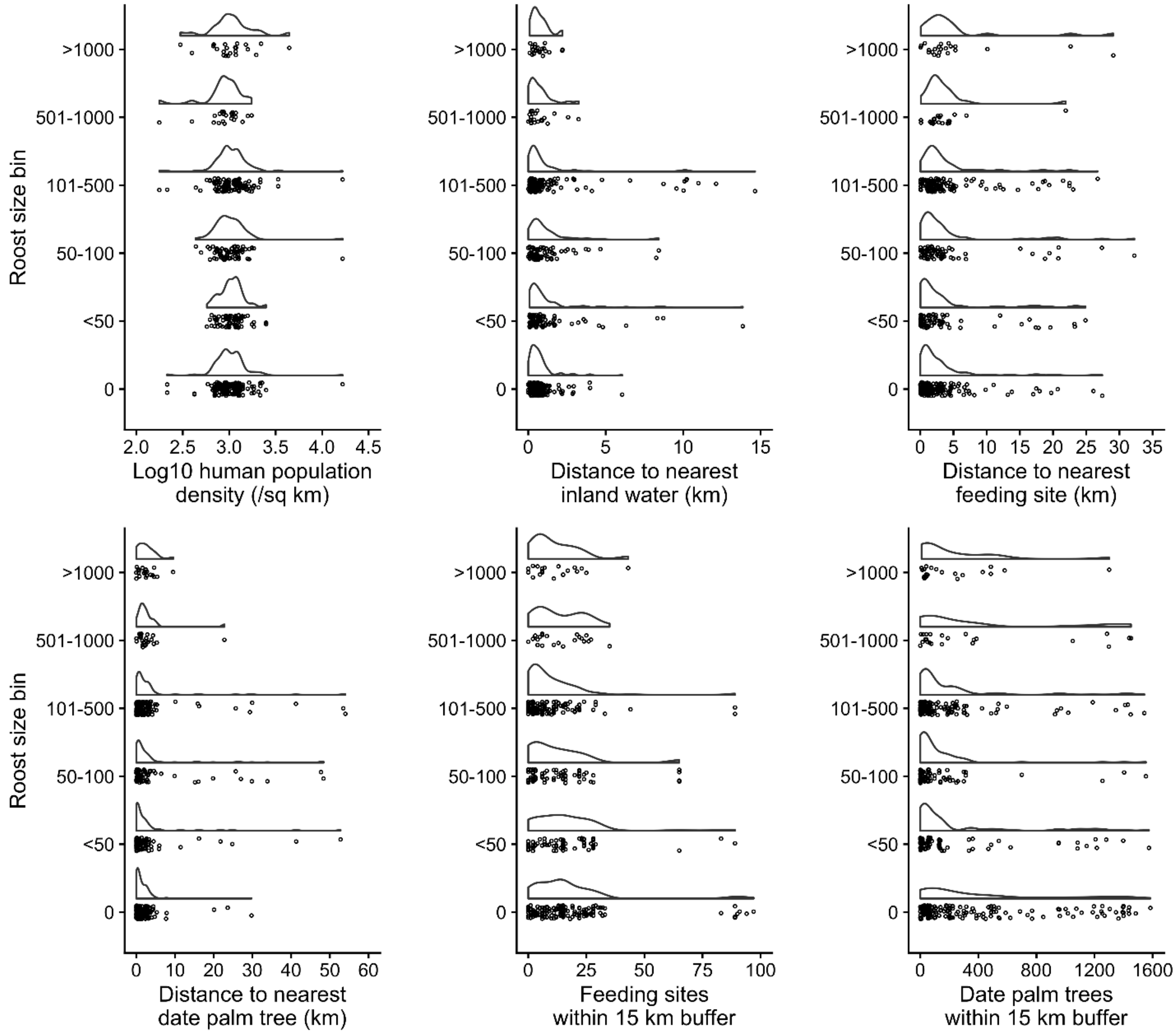


**Figure A8.** Distribution of roost sizes (including unoccupied roost sites) relative to select covariates in Table 2. Raincloud plots show the statistical distribution of variables over individual points.

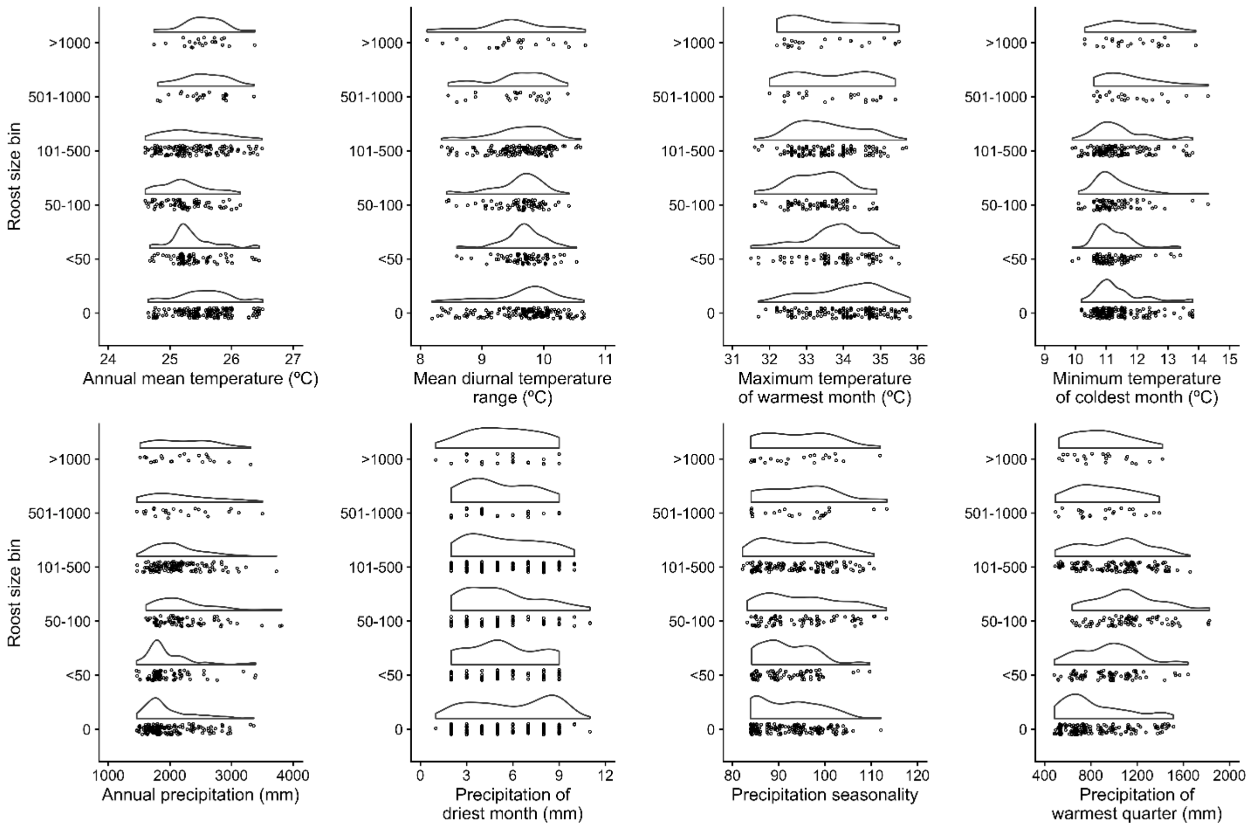


**Figure A9.** Distribution of roost sizes (including unoccupied roost sites) relative to bioclimatic covariates. Raincloud plots show the statistical distribution of variables over individual points.

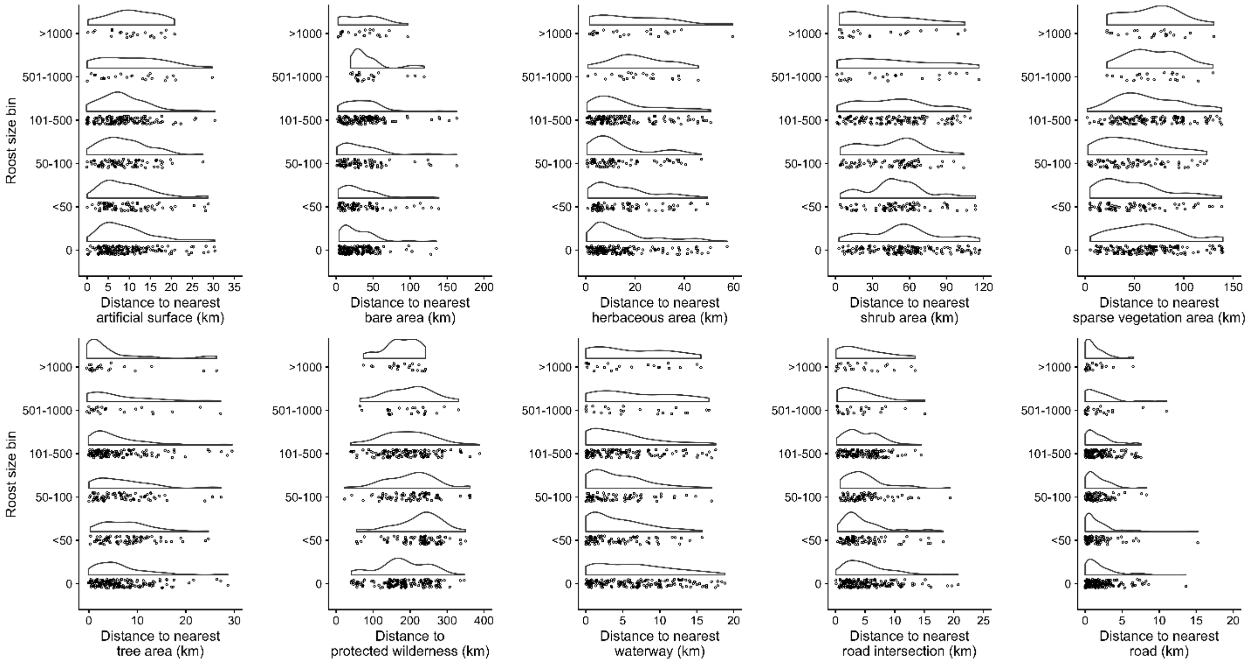


**Figure A10.** Distribution of roost sizes (including unoccupied roost sites) relative to land use covariates. Raincloud plots show the statistical distribution of variables over individual points.

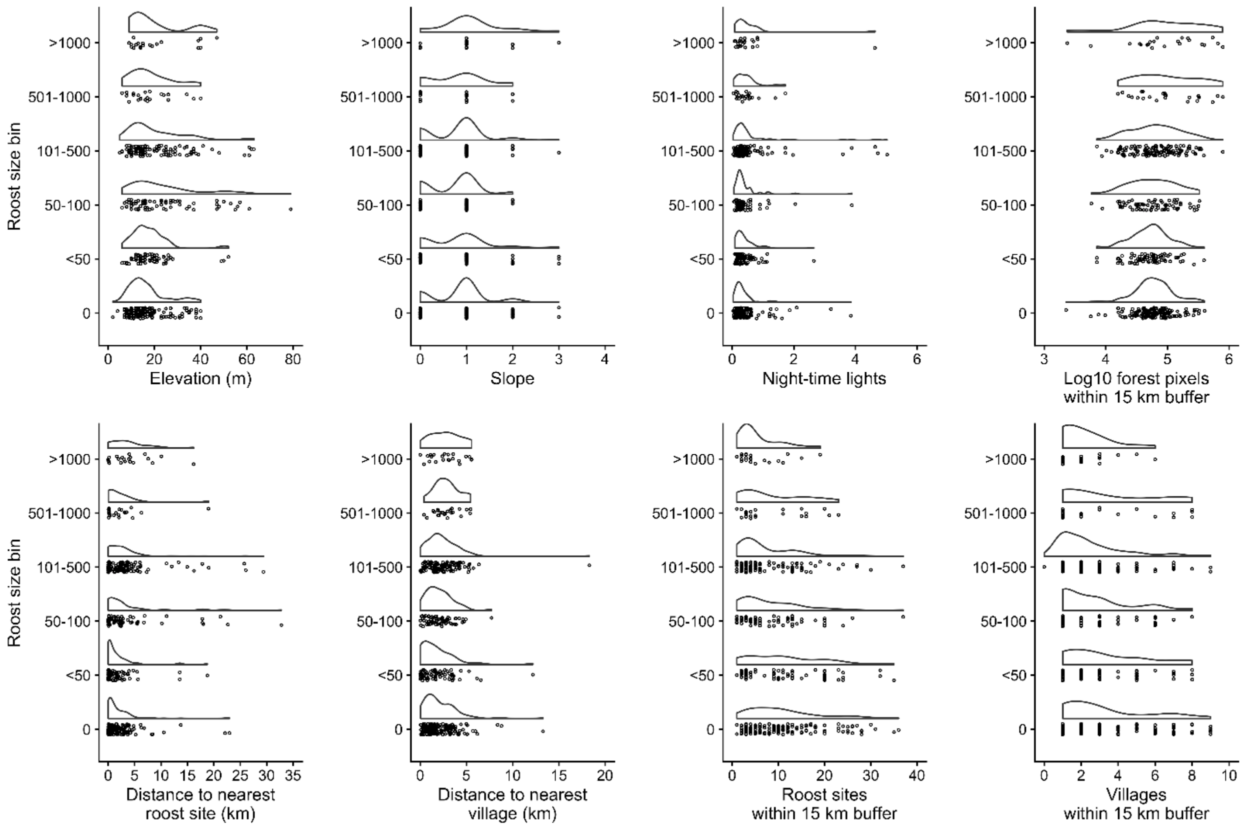


**Figure A11.** Distribution of roost sizes (including unoccupied roost sites) relative to remaining covariates. Raincloud plots show the statistical distribution of variables over individual points.

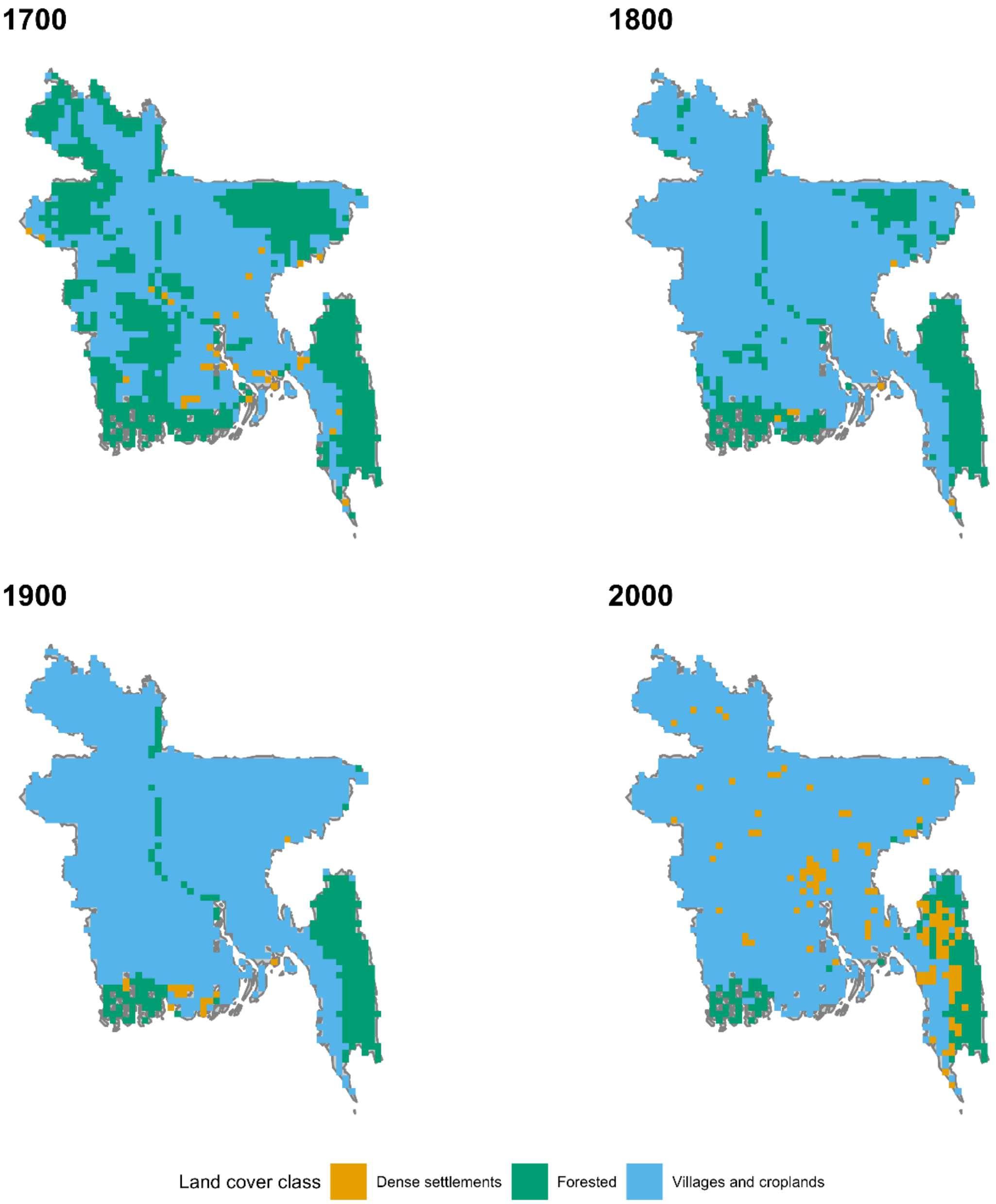


**Figure A12.** Maps of historical change in land cover across Bangladesh. Land cover classes were modified from data in Ellis et al. [6].

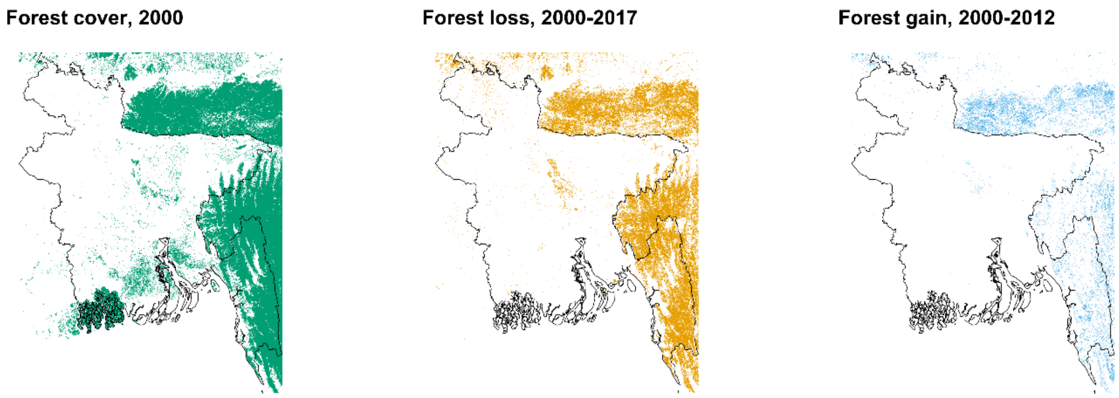


**Figure A13.** Maps of forest cover change in Bangladesh since 2000. Data were drawn from Hansen et al. [5]. Only pixels with forest cover >10% are shown while forest loss and gain within a pixel is binary. Note that forest cover gain only covers the period 2000–2012.
